# Chloroplast activity provides *in vitro* regeneration capability in contrasting cultivars

**DOI:** 10.1101/2022.06.30.498295

**Authors:** Parul Sirohi, Chanderkant Chaudhary, Suchi Baliyan, Ritika Vishnoi, Sumit Kumar Mishra, Reeku Chaudhary, Bhairavnath Waghmode, Anuj Kumar Poonia, Hugo Germain, Debabrata Sircar, Harsh Chauhan

## Abstract

Existence of potent *in vitro* regeneration system is a prerequisite for efficient genetic transformation and functional genomics of crop plants. We know little about why only some cultivars in crop plants are tissue culture friendly. In this study, tissue culture friendly cultivar Golden Promise (GP) and tissue culture resistant DWRB91(D91) were selected as contrasting cultivars to investigate the molecular basis of regeneration efficiency. Multiomics studies involving transcriptomics, proteomics, metabolomics, and biochemical analysis were performed using GP and D91 callus to unravel the regulatory mechanisms. Transcriptomics analysis revealed 1487 differentially expressed genes (DEGs), in which 795 DEGs were upregulated and 692 DEGs were downregulated in the GP-D91 transcriptome. Genes encoding proteins localized in chloroplast and involved in ROS generation were upregulated in the embryogenic calli of GP. Moreover, proteome analysis by LC-MSMS revealed 3062 protein groups and 16989 peptide groups, out of these 1586 protein groups were differentially expressed proteins (DEPs). Eventually, GC-MS based metabolomics analysis also revealed the higher activity of plastids and alterations in key metabolic processes such as sugar metabolism, fatty acid biosynthesis, and secondary metabolism. Higher accumulation of sugars, amino acids and metabolites corresponding to lignin biosynthesis were observed in GP as compared to D91.

**Highlights:** Multi omics analysis revealed chloroplast play crucial role in providing in vitro regeneration capability in contrasting genotypes

## Introduction

Genetic engineering can be used to improve various important traits such as resistance to various biotic and abiotic stresses, to improve the yield and quality of crop plants. For plant genetic engineering, a prerequisite is the availability of an efficient *in vitro* plant regeneration system in the selective crop species regenerating mainly through somatic embryogenesis (SE). *In vitro* regeneration efficiency of plants greatly depends on the formation of embryogenic calli which can be differentiated into new plantlets, and many plant species still have very low somatic embryogenesis efficiency under *in vitro* conditions.

The formation of embryogenic calli is controlled by various intrinsic and extrinsic factors which include the genotype, developmental stage, and explant types in addition to various cultural regimes. Moreover, *in vitro* regeneration and genetic transformation of cereal crops is extraordinarily complex and experimentally challenging. Existing regeneration protocols mainly use immature zygotic embryos as explant and are highly genotype-dependent for most of the crop plants including cereals. Most culture systems for callus formation and plant regeneration were developed for model crop varieties such as ‘Golden Promise’ for barley (Wan and Lemaux, 1994; Tingay et al., 1997; Holme et al., 2006; Kumlehn et al., 2006; Hinchliffe and Harwood, 2019), ‘Bob White and Fielder’ for wheat (Ishida et al., 2015; Pellegrineschi et al., 2002; Hensel et al., 2009; Hayta et al., 2019), ‘TP309 and ’Nipponbare’ for japonica rice (Hiei et al., 1997; Toki, 1997) and ‘IR64 and PB1’ for indica rice (Sahoo et al., 2011), Coker for cotton (Davidonis and Hamilton, 1983; Trolinder and Goodin, 1988; Kumria et al., 2003).

Many plant species have been examined for the characterisation of genes particularly expressed during somatic embryogenesis, including wheat (Chu et al., 2017), cotton (Zeng et al., 2006; Wu et al., 2009; Cheng et al., 2016; Cao et al., 2017), peanut (Rani et al., 2005), carrot (Aleith and Richter, 1991), grape (Gianazza et al., 1992), alfalfa (Domoki et al., 2006), conifer (John and Gerald, 2007; Yakovlev et al., 2016), papaya (Jamaluddin et al., 2017), maize (Salvo et al., 2014) and banana (Enríquez-Valencia et al., 2019). These studies have identified genes that are differentially expressed in SE, including SOMATIC EMBRYOGENESIS RECEPTOR KINASE (SERK) (Schmidt et al., 1997), LEAFY COTYLEDON (LEC) (Curaba et al., 2004), ABSCISIC ACID INSENSITIVE3 (ABI3), FUSCA3 (FUS3) (Freitas et al., 2019), BABY BOOM (BBM) (Boutilier et al., 2002; Maulidiya et al., 2020), WUSCHEL (WUS) (Zuo et al., 2002,), AUXIN RESPONSE FACTOR 19 (ARF19) (Xiao et al., 2020), and LATE EMBRYOGENESIS ABUNDANT (LEA) protein (Wickramasuriya and Dunwell, 2015; Jamaluddin et al., 2017). Although RNA-Seq (RNA Sequencing) analyses have been conducted to investigate genome-wide gene expression in response to somatic embryogenesis (SE) in many plants, the expressions of genes involved along with the key factors determining gene changes and regulations in somatic embryo maturation in cereals have not been previously investigated.

Barley (*Hordeum vulgare* L.) is one of the important cereal crops worldwide and is considered as a model crop for Triticeae tribe for functional genomics studies (Saisho and Takeda, 2011) because of its low genomic size, diploid nature, and available genome sequencing data. The regeneration potential is considered to be highly genotype-dependent and recalcitrant to regeneration under currently available protocols. Hence, the unavailability of robust and reproducible regeneration system in barley is the major limitation for its biotechnological improvement. Genetic transformation of barley through *A. tumefaciens* requires tissue culture dependent regeneration process (Harwood, 2012), which undergo prolonged tissue culture, genotype-dependency and expensive regeneration method. Only few barley cultivars have been reported to produce somatic embryos (SEs) and regenerative plants; and the most responsive cultivar (*cv.*) is Golden Promise (GP) which is a leading cultivar in Europe. In GP, three transformation-amenability loci (*TFA1, TFA2, TFA3*) and one locus in mutant 1460 (*TRA1*) were investigated through genetic mapping that were found to be responsible for Agrobacterium-mediated transformation in barley (Hisano and Sato, 2016; Hisano et al., 2017; Orman-Ligeza et al., 2020).

GP is an excellent cultivar for tissue culture and Agrobacterium mediated transformation but is an old variety prone to many biotic and abiotic stresses. Moreover, it is not suitable for climatic conditions outside temperate zones, whereas most new commercial barley cultivars are not tissue culture friendly and show recalcitrant embryogenesis from explant in genetic transformation, which hinders its application in transgenic breeding. In the present study, we analyzed the transcriptomic profiles and gene regulation changes in the somatic embryo maturation process of two contrasting cultivars of barley (GP and DWRB91 (D91)). The study also aimed to determine a possible relationship between somatic embryogenesis and some metabolic contents in embryogenic and non-embryogenic calluses of GP and D91. The results from this study will help to improve our understandings of the molecular mechanisms in the somatic embryogenesis of barley along with other cereal crops and will offer a fresh insight on how genetic modification can increase the effectiveness of somatic embryogenesis.

## Material & Method

### Plant material and somatic embryogenesis

Seeds of barley cultivars were provided by ICAR-National Bureau of Plant Genetic Resources New Delhi. Mature seeds were sterilized with 4% sodium hypochlorite solution for 20 min and then rinsed with sterile water 4-5 times. For induction of embryogenic callus, mature embryos were dissected from seeds with a sterile surgical blade and put with scutellum side up on the barley callus induction media (BCIM) containing 4.4 g L^−1^ Murashige and Skoog (MS) plant salt base (Duchefa M0231), 5 µM CuSO4, 1 mgL^−1^ Thiamine HCl, 0.69 gL^−1^ L-Proline, 2.0 mgL^−1^ 2,4-D, 30 gL^−1^ Maltose.H20, 1 gL^−1^ Casein hydrolysate, 0.25 gL^−1^ Myo-inositol, 3 gL^−1^ Phytagel, pH 5.8 (Hensel et al., 2009). Mature embryos were cultured at 22 °C under a 16 h light/8 h dark photoperiod for 28 days. After 28 days, calli were transferred to regeneration medium (BRM) for shoot formation. BRM consisted of 4.4 g L^−1^ Murashige and Skoog (MS) plant salt base (Duchefa M0231),146 mgL^−1^ L-Glutamine, 0.225 mgL^−1^ 6-BAP, 36 gL^−1^ Maltose·H2O, 3 gL^−1^ Phytagel, pH 5.8 (Hensel et al., 2009). The callus derived from mature embryos of GP and D91 were harvested at different time intervals as specified for further analysis. Similarly, immature embryos of respective cultivars were isolated from developing seeds at 12-14 days after anthesis, from plants grown in green house.

### Scanning Electron Microscopy (SEM)

4-weeks-old calli (28 days on BCIM in dark + 3 days on BRM under indirect light) of GP and D91 were chosen in order to examine the callus’ outer surface under a scanning electron microscope (Hitachi, Tm 30000, Tokyo, Japan). The samples were prepared as described by Bomblies et al. (2008). Briefly, the calli were placed in fixative of formaldehyde-acetic acid (FAA; 3.7% v/v formaldehyde, 5% v/v acetic acid, 70% v/v ethanol) solution with brief application of vacuum to remove air bubbles and then incubated for 24 hr at 4℃. The fixative was replaced with 1% OsO4 (v/v) to cover the samples and incubated overnight at 4°C. Fixed tissue samples were rinsed three times in 25 mM sodium phosphate (pH 7.2), followed by dehydrating the tissue with an ethanol series (30%, 50%, 70%, 95%, and twice in 100% ethanol) at room temperature for at least 15-30 min each. The calli were then dried using critical point drying for one hour at 35 ℃ (CPD: Leica). Chromium coating was done on dried calli followed by observation in SEM.

### Relative water content

A measure that reflects cellular osmotic stress is relative water content (RWC). Callus samples of known fresh weight were kept in milliQ water for 12 hours and then weighed for their full turgid weight. To determine the callus dry weight, the samples were subsequently dried for 48 hrs in an oven set at 65°C. They were then reweighed in order to calculate the difference between their initial and final masses. RWC was calculated as [(fresh weight – dry weight)/(turgid weight – dry weight) × 100] (Pieczynski et al., 2013.).

### Transcriptome profiling using RNA-Seq

Total RNA was isolated from 31-day-old calli sample (28 days BCIM dark + 3 days BRM indirect light) of GP and D91 using the RNeasy plant RNA isolation kit with on column DNase digestion, as per manufacture’s instruction (Qiagen, Hilden, Germany; Cat No./ID: 74904 and 79254). Three biological replicates were used for each cultivar. For RNA-Seq using the Illumina HiSeq technology with paired-end read length of 101 bp, high-quality RNA samples were sent to Bionivid Technology Private Limited (Bengaluru, India). All reads were subjected to quality control using the NGSQC Toolkit (Patel and Jain 2012), and reads with Phred score> 20 were chosen for downstream analysis. Reference genome of barley was downloaded from Ensembl Plants database (http://ftp.ensemblgenomes.org/pub/plants/release-53/fasta/hordeum_vulgare/) for alignment and mapping of reads using STAR pipeline (Dobin et al., 2013; https://github.com/alexdobin/STAR) and Trinity platform (genome guided mode) (Hass et al., 2013; https://github.com/trinityrnaseq/trinityrnaseq). DESeq2 (Mi et al., 2014; https://bioconductor.org/packages/release/bioc/html/DESeq2.html) was used to analyse differential gene expression, and the RSEM (RNA-Seq by Expectation-Maximization) approach was used to estimate abundance (Li and Dewey, 2011). Transcripts having log2 fold change ≥ 2 were considered as DEGs. Functional annotation of the transcriptome was performed using the Trinotate pipeline (https://github.com/Trinotate/Trinotate.github.io/blob/master/index.asciidoc) employing uniport-Swissprot (https://ftp.uniprot.org/pub/databases/uniprot/current_release/knowledgebase/complete/) database for Blastx and Blastp searches. Cytoscape (v 3.9.1) plug-in bingo (Maere et al., 2005) was used to perform GO (Gene Ontology) enrichment analysis. Enriched GO terms were identified separately for upregulated and downregulated sets of genes in GP as compared to D91. Obtained enriched terms were plotted using the web server Revigo (http://revigo.irb.hr/; Supek et al., 2011). Heatmaps were constructed using ggplots (Warnes et al., 2020) and RColorBrewer (Neuwirth, 2014) packages in R.

### Gene validation through quantitative RT-PCR

RNA was isolated from GP and D91 callus tissues selected at different time intervals during callus induction (3 days, 7 days, 14 days, 21 days, 28 days, 31 days (28 days on CIM + 3 days on BRM)) and was used for qRT-PCR. Different time intervals for calli was taken to analyze the expression of selected genes during the entire process of callus induction, formation of somatic embryos followed by maturation. RNA concentration was determined using a NanoDrop OneC Microvolume UV-Vis Spectrophotometer (Thermo Scientific, USA), and the quality of the RNA was assessed by running on 1 % agarose gel in MOPS formaldehyde buffer. In short, the iScript^TM^ cDNA synthesis kit (Bio-Rad, Cat. No. 1708891) was used to synthesize cDNA from 1 µg of RNA in accordance with the manufacturer’s instructions. qRT-PCR was used for the expression analysis, with the QuantStudio 3 system and PowerUp SYBR Green Master Mix (Applied Biosystems, Thermo Fisher Scientific, Cat. No. A25741). snoR14 was used as internal standard and the fold change in gene expression was calculated using the formula 2^-ΔΔCt^ (Risk et al., 2013). Three biological replicates and three technical replicates were used for qRT-PCR reactions. Primers from selected genes were designed using coding DNA sequences (CDS) and Primer express 3.0 software (Thermo Fisher Scientific). Supplementary Table S1 contains a full list of primers and the gene IDs for the respective barley genes.

### Protein extraction and proteome analysis using LC-MS/MS

Protein was extracted using phenol method from callus tissues of *cv.* GP and *cv.* D91. Approximately 1 g of tissue was crushed using mortar-pestle in liquid nitrogen and tissue powder was homogenized using 3 ml of extraction buffer (500 mM Tris-HCl, 700 mM sucrose,50 mM EDTA, 100 mM KCl; pH 8.0; 2% 2-mercaptoethanol, and 1 mM phenyl methyl sulfonyl fluoride (PMSF)) followed by vortexing and incubation for 10 min on ice. Afterward, an equal volume of Tris-buffered phenol (pH 8.0) was added to the extract, and centrifuged at 12000g for 10 min at 4°C. The upper phenol phase was transferred to a fresh tube and the phenolic phase was re-extracted with 3 ml of extraction buffer. Protein was precipitated using 4 volume of 0.1M ammonium acetate (dissolved in cold methanol) for 4 hr at -20°C. Proteins were finally pelleted by centrifugation at 5500g for 10 min and final pellet was dissolved in lysis buffer (5% SDS and 0.1M Tris (pH 8.8). Protein was quantified using Bradford reagent (Sigma-Aldrich) and samples were stored at -80°C until further use.

TCEP (tris(2-carboxyethyl) phosphine) was used to reduce a protein sample of 50 µg, which was then further alkylated with 50 mM iodoacetamide before being digested with trypsin (1:50 trypsin/lysate ratio) for 16 hr at 37°C. The digests were purified with a C18 silica cartridge, followed by drying using speed vac. The dried pellet was resuspended in buffer A (2% acetonitrile, 0.1% formic acid). MS analysis was performed using *EASY-nLC^TM^ 1200 system* (Thermo Fisher Scientific) coupled to *Q Exactive^TM^ plus* equipped with nanoelectrospray ion source (Thermo Fisher Scientific). 1 μg sample was loaded on 50 cm C18 column, 3.0μm Easy-spray column (Thermo Fisher Scientific) and peptides were eluted with a 0–40% gradient of buffer B (80% acetonitrile, 0.1% formic acid) at a flow rate of 300 nl/min. LC gradients were run for 100 minutes. MS1 spectra were acquired in the *Orbitrap* at 70k resolution. All charge states for a specific precursor were excluded using dynamic exclusion for 10 seconds while MS2 spectra were collected at a resolution of 17500.

### Data acquisition and analysis

Raw files were analysed with Proteome Discoverer (v2.2) (Thermo Fisher Scientific) against the Uniprot Knowledge Base (UniprotKB) *Hordeum vulgare* proteome database (https://www.uniprot.org/proteomes/UP000011116, taxon ID: 112509, 1899799 entries). SEQUEST (Eng et al., 1994) and MS Amanda (Dorfer et al., 2014) searches were performed using the precursor and fragment mass tolerances at 10 ppm and 0.02 Da, respectively. The enzyme specificity was set for trypsin/P (cleavage at the C terminus of “K/R: unless followed by “P”) to generate peptides with maximum missed cleavages value of 2. Fixed modification as carbamidomethyl on cysteine (C) and variable modification as oxidation of methionine (M) were used for database search. Both peptide spectrum match and protein false discovery rate (FDR) were set to 0.01 and filtered values were log2 standardized followed by imputation of missing values on the basis of normal distribution. T-test was performed on normalised values and significance was calculated using Benjamini-Hochberg adjusted p-value. Graphical representations were plotted using the significant proteins (DAPs; Differentially Abundant Proteins) in R. To functionally group proteins we also used BLASTP searches locally against the uniport-swissprot (https://ftp.uniprot.org/pub/databases/uniprot/current_release/knowledgebase/complete/uniprot_sprot.fasta.gz) database.

### Construction of binary vectors for subcellular localization

To check the subcellular localization of targeted genes, Gene::GFP fusion cassettes driven by the constitutive *35S CaMV* promoter were constructed. The ORF of selected DEGs without their stop codon were PCR-amplified using primer pairs (Supplementary Table S2) and cloned into entry vector *pDONR221* (Thermo Fisher Scientific, Waltham, MA, USA) and subsequently into destination vector *pB7FWG2* (Karimi et al., 2002) by Gateway™ cloning [Invitrogen™ Gateway™ BP Clonase™ II Enzyme mix (Cat No.- 11789020) and Invitrogen™ Gateway™ LR Clonase™ II Enzyme mix (Cat No.- 11791020) to construct *pB7FWG2-35S::Gene:GFP*. The cloned sequences were verified by sequencing.

### Transient expression in *Nicotiana benthamiana* and confocal laser scanning microscopy

*A. tumefaciens* was infliltrated into the *Nicotiana benthamiana* leaves abaxially using the method described by Li (2011). The vector *pB7FWG2-35S::Gene:GFP* was transformed into *A. tumefaciens* GV3101. For agroinfiltration, *A. tumefaciens* GV3101 harbouring the binary vector was cultured overnight in LB-broth media containing 50 mg/L of spectinomycin, 30 mg/L of rifampicin, 30 mg/L gentamycin, to the stationary phase in the shaking incubator with 250 rpm at 28 °C. Bacterial cells were harvested by centrifugation and resuspended in the infiltration medium (10 mM MgCl2, 100 μM acetosyringone) to a final OD600 of 0.5. The suspension was infiltrated into the abaxial surface of 4-weeks-old leaves of *N. benthamiana* with a 1 mL needleless syringe (Wroblewski et al., 2005). *N. benthamiana* plants were kept at 25℃ for transient gene expression post agroinfiltration. A confocal laser scanning microscope (TCS SP8, Leica Microsystems, Wetzlar, Germany) was used to examine protein localisation 24–48 hours after infiltration. GFP was excited at 488 nm and the fluorescence detected between 498–525 nm. Autofluorescence of the chloroplasts was detected at 680–700 nm.

### Histochemical detection and quantification of superoxide anions

28-days calli (BCIM) and 31-days calli (28 days BCIM + 3 days BRM) were vacuum infiltrated in a desiccator in a 5 mL nitroblue tetrazolium (NBT) staining solution containing 0·1% (w/v) NBT in 10 mM MES, pH 6.5, 10 mM sodium azide, 50 mM potassium phosphate, pH 6.4 (Wohlgemuth et al., 2002). The calli were incubated for 24 hrs in dark at room temperature. Formazon precipitated in NBT stained calli was selectively removed with chloroform for quantitative measurement. After the organic phase was dried, formazan residue was dissolved in dimethylsulfoxide-potassium hydroxide (DMSO-KOH) and measured using spectrophotometry (Grellet Bournonville and Díaz-Ricci, 2011).

### Metabolite sample preparation

Metabolite extraction for GC-MS analysis was done as per the protocol described by Sarkate et al. (2018). Briefly, 31-day callus tissue (∼500mg) of GP and D91 was collected and dried in hot air oven at 60 °C for 4 h. For metabolite extraction, dried callus was crushed in liquid nitrogen to produce fine powdered samples. 500 mg of powdered materials were placed in a 1.5 mL micro- centrifuge tube together with 1 ml of a pre-cooled extraction mixture containing methanol, water, and chloroform in the ratio of 2.5:1:1 (v/v/v) followed by vigorous vortexing for 2 min at room temperature. As an internal reference to determine extraction efficiency, 50 µL of 2-Phenylphenol (2 mg/mL methanol stock) was added to the extraction mixture and centrifuged at 14000 x g for 5 min. The supernatant (0.8 mL) was collected into a new 1.5 mL micro-centrifuge tube and then 0.4 mL of water was added, the mixture was vortexed for 10 sec followed by centrifugation at 14000 x g for 5 min. The polar top solvent partition phase (methanol/water) was transferred to a fresh micro-centrifuge tube, dried out in a vacuum concentrator (Eppendorf Concentrator plus™, Eppendorf; USA) at 20 °C for 2 hr, and then freeze dried for 12 hr in a lyophilizer. For GC-MS analysis, dried material was ultimately doubly derivatized (Lisec et al., 2006). In the first derivatization, 40 μL of methoxyamine hydrochloride (stock solution: 20 mg/ml in pyridine) was added to the dried sample and incubated at 37 °C for 2 hr. Using 50 µL of N-methyl-N- (trimethylsilyl)-trifluoroacetamide (MSTFA) at 37 °C for 60 min, the second derivatization was carried out. Subsequently, second derivatization was carried out by adding 80 μL of N-methyl-N- (trimethylsilyl)-trifluoroacetamide (MSTFA) and then incubation at 37 °C for 30 min. As a negative control, a derivatization process was prepared using an empty tube without a sample. The mixture was centrifuged at 14000 g for 10 min after derivatization was completed. A fresh GC glass vial was used to transfer the resultant supernatant, and it was then promptly analysed by GC-MS to look for primary and secondary metabolites.

### GC-MS analysis

The GC-MS studies were conducted using an Agilent gas chromatograph (7890B) and mass detector (5977B) (Agilent Technologies, CA, USA). On an HP-5 MS column (5 % phenyl methyl polysiloxane; 30 m length x 0.25 mm i.d. x 0.25 µm thickness; Agilent technologies, CA, USA), metabolites were separated. Helium was used as the carrier gas with a 3 µL injection volume and a 1 mL min^-1^ flow rate. The oven temperature was initially set at 70 °C for 5 min, then increased from 70 °C to 270 °C at a rate of 5 °C min^-1^, then held at 270 °C for 1 min, and then again increased from 270 °C to 300 °C at a rate of 10 °C min^-1^, kept at 300 °C for 15 min. The GC operation conditions were set as standard with 5:1 split mode, injector temperature 250 °C and transfer line temperature 200 °C. Temperatures for the MS source and MS quad were set to 230 °C and 150 °C, respectively. The mass detector was run in electron ionisation mode at 70 eV with a scanning range of m/z 30-750 and a scanning speed of 0.34 s. Each sample was replicated three times. The method outlined in Waghmode et al., (2021) was used to identify the metabolites, and each one was identified by matching its mass spectral data to the National Institute of Standards & Technology library (NIST 17.L) database.

### Metabolite data pre-processing and statistical analysis

WsearchPro’s tools were used to deconvolute raw GC-MS data files collected from Agilent ChemStationTM software using Automated Mass Spectral Deconvolution and Identification System (AMDIS) (www.wsearch.com.au). Before uploading to Metaboanalyst 3.0 (http://www.metaboanalyst.ca), the resulting metabolite data files from GP and D91 were consolidated into a single file and converted to.csv (comma separated values) format. Data were normalized using area of internal standard (2-phenylphenol). Data were then log converted using Pareto scaling (mean centred and divided by the square root of each variable’s standard deviation) before statistical analysis. One-way analysis of variance (ANOVA) was performed using the Statistical Package for the Social Sciences (SPSS) software to determine significant differences in metabolite levels. Tukey’s significant-difference test was then performed with a statistical significance level of p < 0.05. The metabolic profiles of the GP and D91 samples, which were depicted as 2D score plots, were distinguished using partial least squares discriminant analysis (PLS-DA) and principal component analyses (PCA). To differentiate between samples, key characteristics from the PLS-DA data were scored according to their variable importance in projection (VIP). Heatmaps were created using Metaboanalyst 3.0 by using Pearson distance measure and the Ward clustering algorithm. Finally, using the pathway analysis tool in Metaboanalyst 3.0, a simplified metabolic pathway interaction map was manually generated using data from the Kyoto Encyclopedia of Genes and Genomes (KEGG) database.

### Lignin estimation and phloroglucinol staining

Lignin estimation was done in 4-weeks-old calli as per the method described by Dampanaboina et al. (2021). For lignin staining, method described by Mitra and Loque (2014) was used. The phloroglucinol-HCl (Ph-HCl) or Wiesner stain was prepared freshly by dissolving 0.3 g in 10 ml of 100% ethanol, which was then combined with two volumes of concentrated HCl (37 N) to make a 3% solution. Calli was transferred in a scintillation vial and Ph-HCl solution was added to completely dip the calli for staining.

## Results

With an initial aim to identify a suitable Indian commercial cultivar showing *in vitro* regeneration response equivalent to GP, we checked 13 genotypes of Indian barley. We found that all the genotypes showed comparable callusing on 2,4-D supplemented media in both immature and mature embryo explants (Supplementary Fig S1A). However, when it come to regeneration response, only DWRB73, DWRB52 and RD2668 showed comparable regeneration response in terms of both percent regenerating calli as well as average shoots per callus regenerated (Supplementary Fig S1B). Also, we found that cultivar namely DWRB91 and DWRB92 never produced regenerated shoots in both immature and mature embryo derived calli, except some occasional greening upon longer time period (8-12 weeks on regeneration media). For further experiments, we choose GP and DWRB91 (D91) as contrasting genotypes.

### Callus morphology and scanning electron microscopy

Somatic embryogenesis, i*n vitro* regeneration and transformation efficiencies are highly genotype dependent in crop plants. In the present study, mature embryos of two contrasting barley genotypes *vi*z. GP and D91 were used as explants for callus induction to find more about molecular basis of *in vitro* regeneration. Both the cultivars showed callus initiation within 48 hr from the scutellum on MS media supplemented with 2,4-D. The calli were visualized for somatic embryogenesis for 4-weeks and there is clear distinction between the callus morphology between these two cultivars while GP developed pale yellow or creamy compact calli (dense and granular) and D91 developed soft friable, white watery calli (loosely arranged) after 4-weeks of culture (Fig. 1A). Greening along with shoot formation in almost 90% of calli (4-weeks-old) of GP was observed after transferring to regeneration media, whereas in case of D91 only little greening was observed with no shoot formation at all (Fig. 1B). The results show high embryogenic potential of GP as compared to D91.

**Fig. 1.**
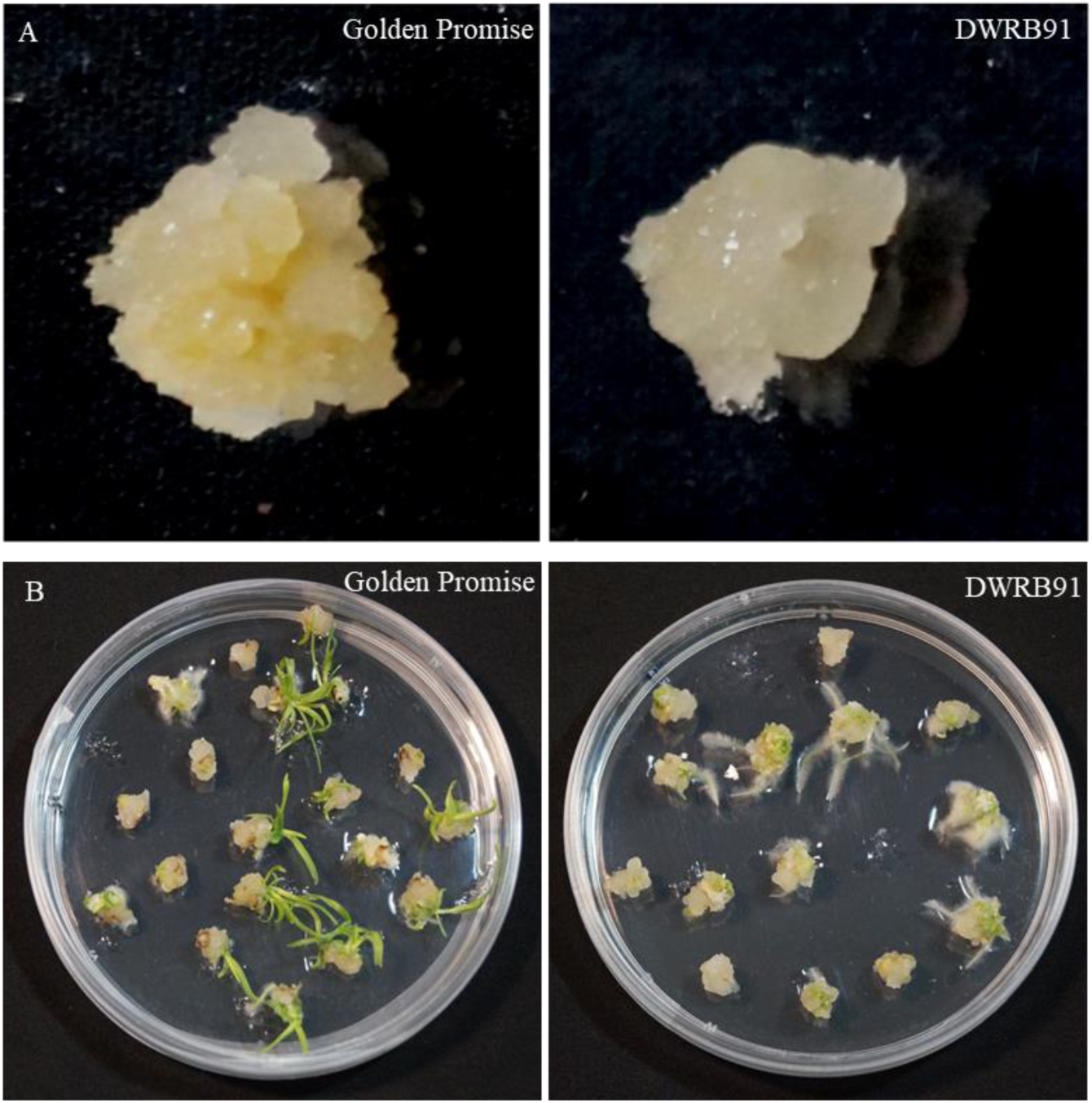
Morphological analysis of Golden Promise and DWRB91 callus. (A) Pale yellow, compact embryogenic calli of Golden Promise vs white, watery non-embryogenic calli of DWRB91. (B). Callus of Golden Promise and DWRB91 showing difference in regeneration capacity on BRM media.

The appearance of the callus surface of embryogenic and non-embryogenic calli was examined using scanning electron microscopy (SEM). The 4-weeks-old calli of GP and D91 were collected for SEM analysis. Golden Promise calli were different morphologically, consisting of distinct compact, nodular, and spherical cells that were securely held together and appeared to arrange globular embryos in discrete clusters, according to SEM analysis (Fig. 2A). However, D91 calli showed tubular cells which were elongated and loosely held on the surface (Fig. 2B). The results of SEM observation demonstrated that GP calli have higher embryogenic potential than D91 calli.

**Fig. 2.**
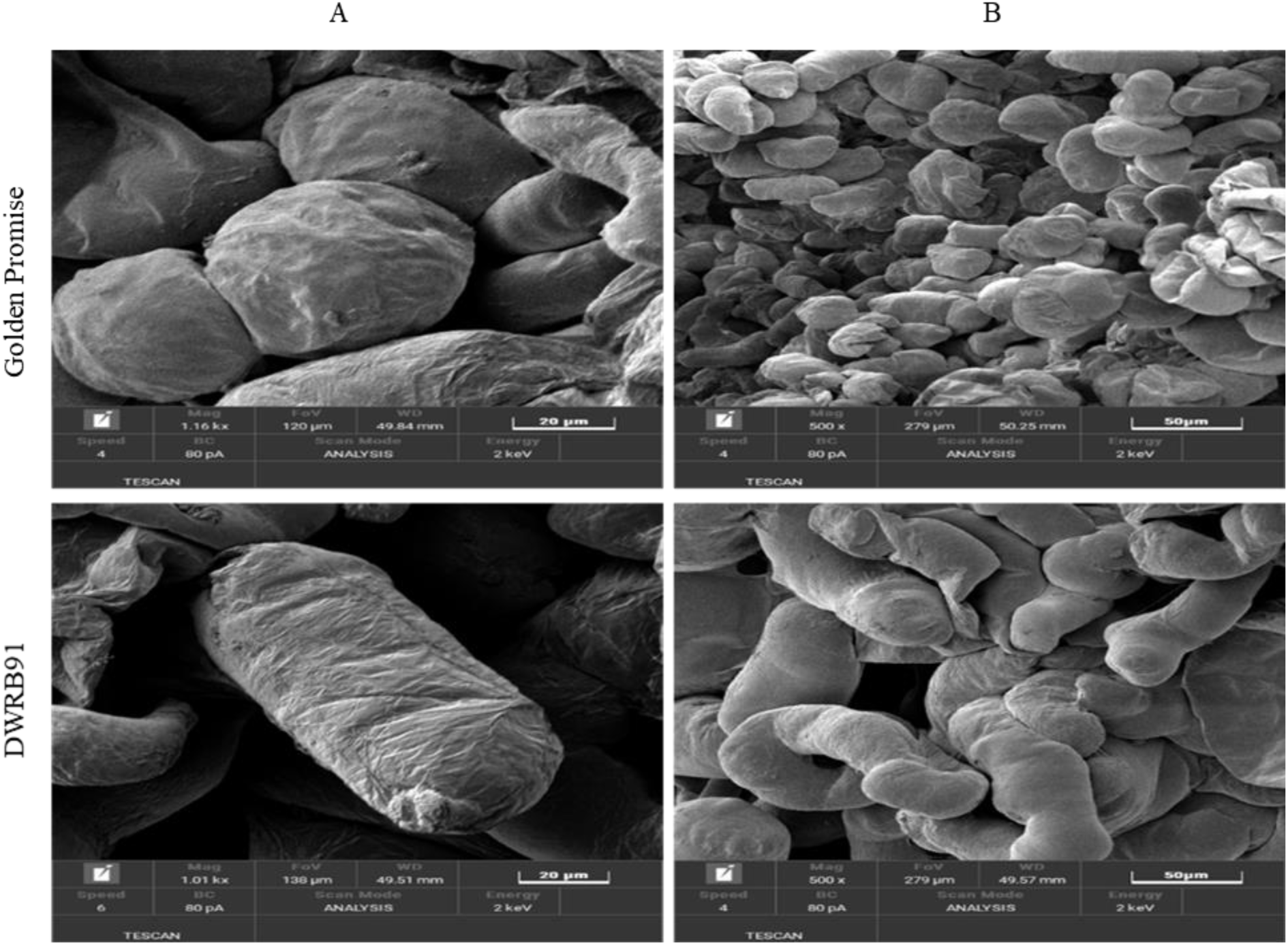
Scanning electron microscopic observation of 4-weeks callus of Golden Promise and DWRB91. Bar A =20 µm, B = 50 µm

Relative water content for calli of both GP and D91 was calculated as the ratio of callus water content determined by the absolute difference of callus fresh weight and dry weight by drying at 70 ℃ to the fresh weight of callus. Significant differences in the relative water content of GP and D91 calli was observed (Fig. 3). The results show significantly higher relative water content in D91 callus (95 %) as compared to GP callus (78 %), which is in line with the visual appearance of calli from both these genotypes.

**Fig. 3.**
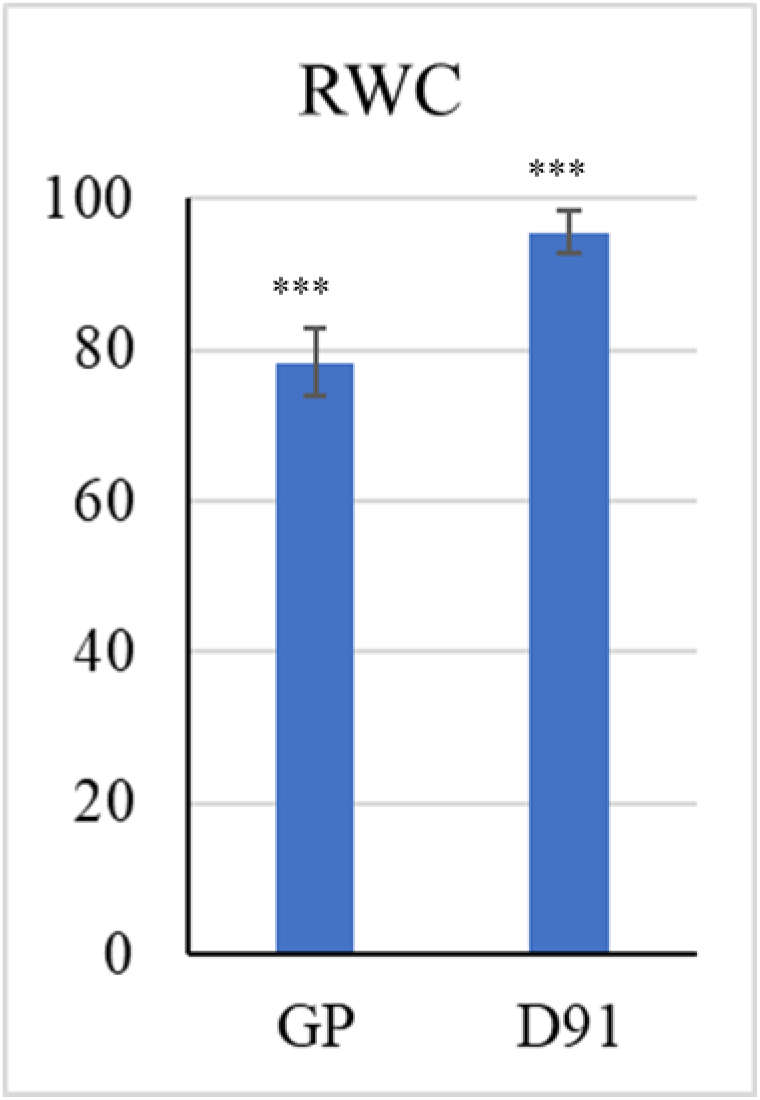
Relative water content of Golden Promise and DWRB91 calli. Data represented as the mean values +- SD of three biological replicates. *** represents the P value significance < 0.001

### GP-D91 transcriptome profile

High-throughput RNA-Seq was performed for 4-weeks-old calli of barley *cv.* GP and *cv.* D91 to understand the molecular mechanisms of somatic embryogenesis, which is a major challenge during *in vitro* regeneration and genetic transformation. A total of 112 million raw reads were generated, with approximately 51 million and 58 million high-quality reads for GP and D91, respectively (Supplementary Table S3).

A total of 1487 DEGs were identified, out of these, 795 DEGs were upregulated and 692 DEGs were downregulated in the embryogenic callus produced by GP as compared to non-embryogenic callus of D91 (Table 1). Gene Ontology (GO) enrichment analysis revealed biological processes such as “photosynthesis”, “photosystem II stabilization”, “regulation of photosynthesis”, “photosystem II oxygen evolving complex assembly”, “developmental growth”, “meristem determinacy”, “suspensor development”, “embryonic pattern specification”, “carbohydrate biosynthetic process”, “glycoside metabolic process”, “chlorophyll metabolic process”, and “actin polymerization” were found to be significantly enriched in the upregulated DEGs of *cv.* GP (Fig. 4A and Table 2A) whereas terms related to “root morphogenesis”, “regulation of ARF protein signal transduction”, “phosphate ion homeostasis”, “amyloplast organization”, “plant-type cell wall loosening”, and “fucose metabolic process” were enriched in the downregulated DEGs of *cv.* GP (Fig. 4B and Table2B). Similarly, cellular component and molecular function GO terms were identified for the DEGs. Cellular component terms such as “cytoskeleton”, “spindle”, “chloroplast”, “plastid”, “chloroplast photosystem II”, and “photosystem” were enriched in the upregulated DEGs (Fig. 4A and Table 2A) whereas in the molecular function category, “sucrose synthase activity”, magnesium-protoporphyrin IX monomethyl ester cyclase activity, and “oxygen evolving activity”, terms were enriched in upregulated DEGs of *cv.* Golden Promise (Fig. 4A and Table 2A). Taken together, DEGs and GO enrichment analysis suggested the upregulation of genes related to primary metabolisms such as photosynthesis, chlorophyll biosynthesis, and plastid formation in the embryogenic callus of *Hordeum vulgare cv.* GP unlike the non-embryogenic callus of *cv.* D91.

**Fig. 4.**
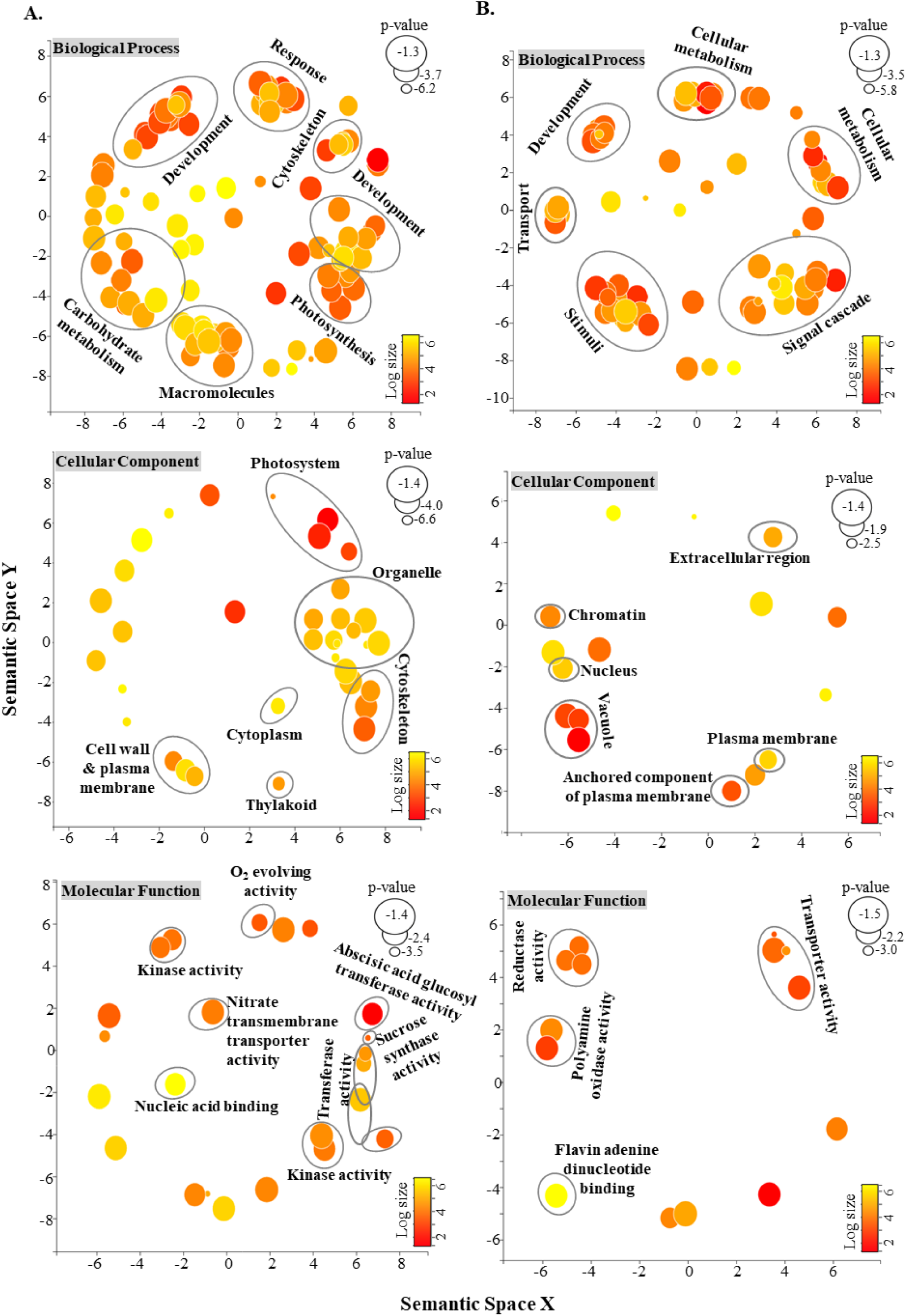
GO enrichment analysis. Scatterplot showing the enriched biological processes, molecular function, cellular component GO categories for DEGs in the calli of barley *cvs.* (A)

**Table 1.**
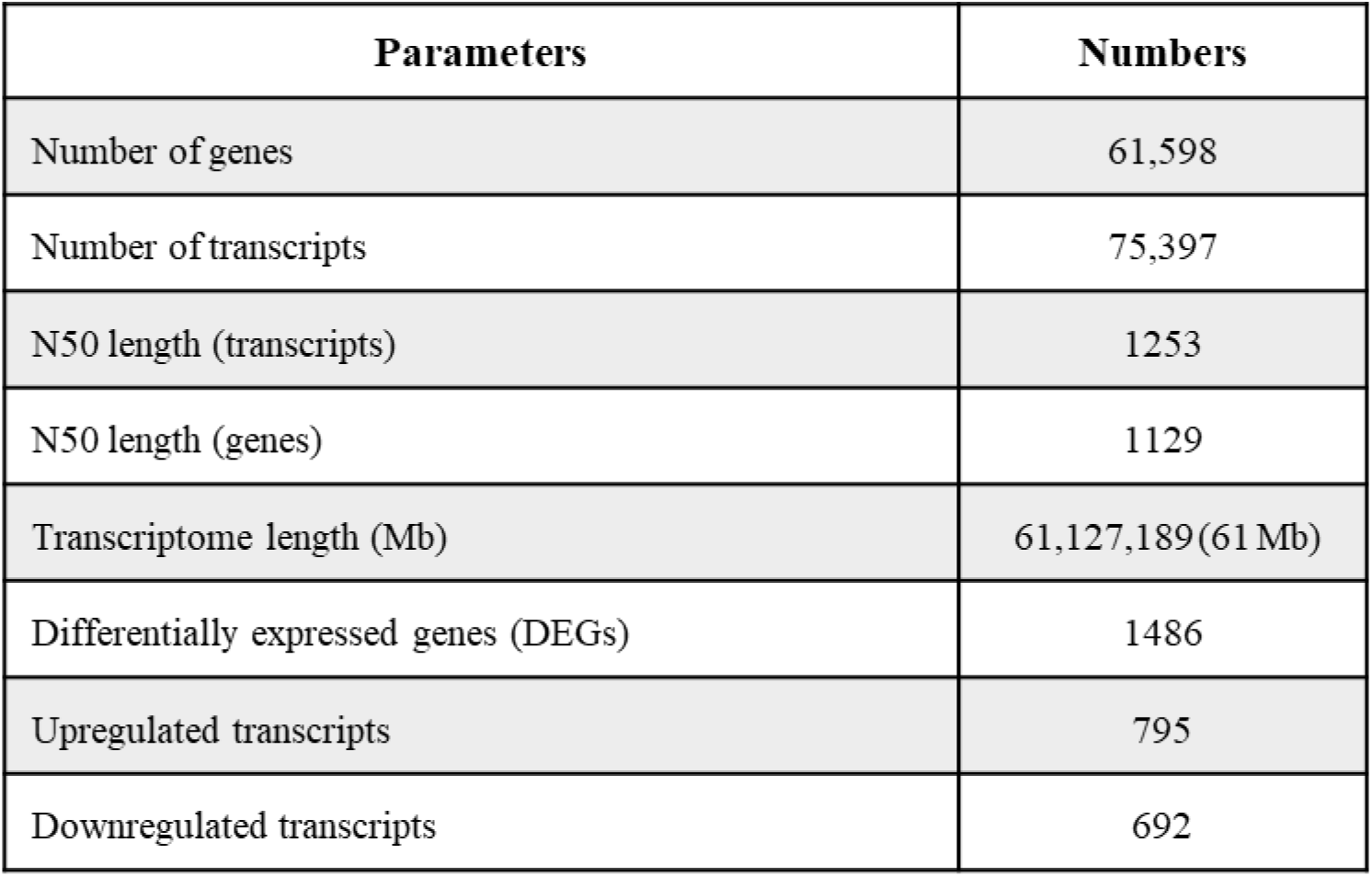
Summary of genome guided assembly of barley calli (*Hordeum vulgare cv.* GP and *cv*. D91) transcriptome.

| Parameters | Numbers |
| --- | --- |
| Number of genes | 61,598 |
| Number of transcripts | 75,397 |
| N50 length (transcripts) | 1253 |
| N50 length (genes) | 1129 |
| Transcriptome length (Mb) | 61,127,189(61 Mb) |
| Differentially expressed genes (DEGs) | 1486 |
| Upregulated transcripts | 795 |
| Downregulated transcripts | 692 |

**Table 2A.**
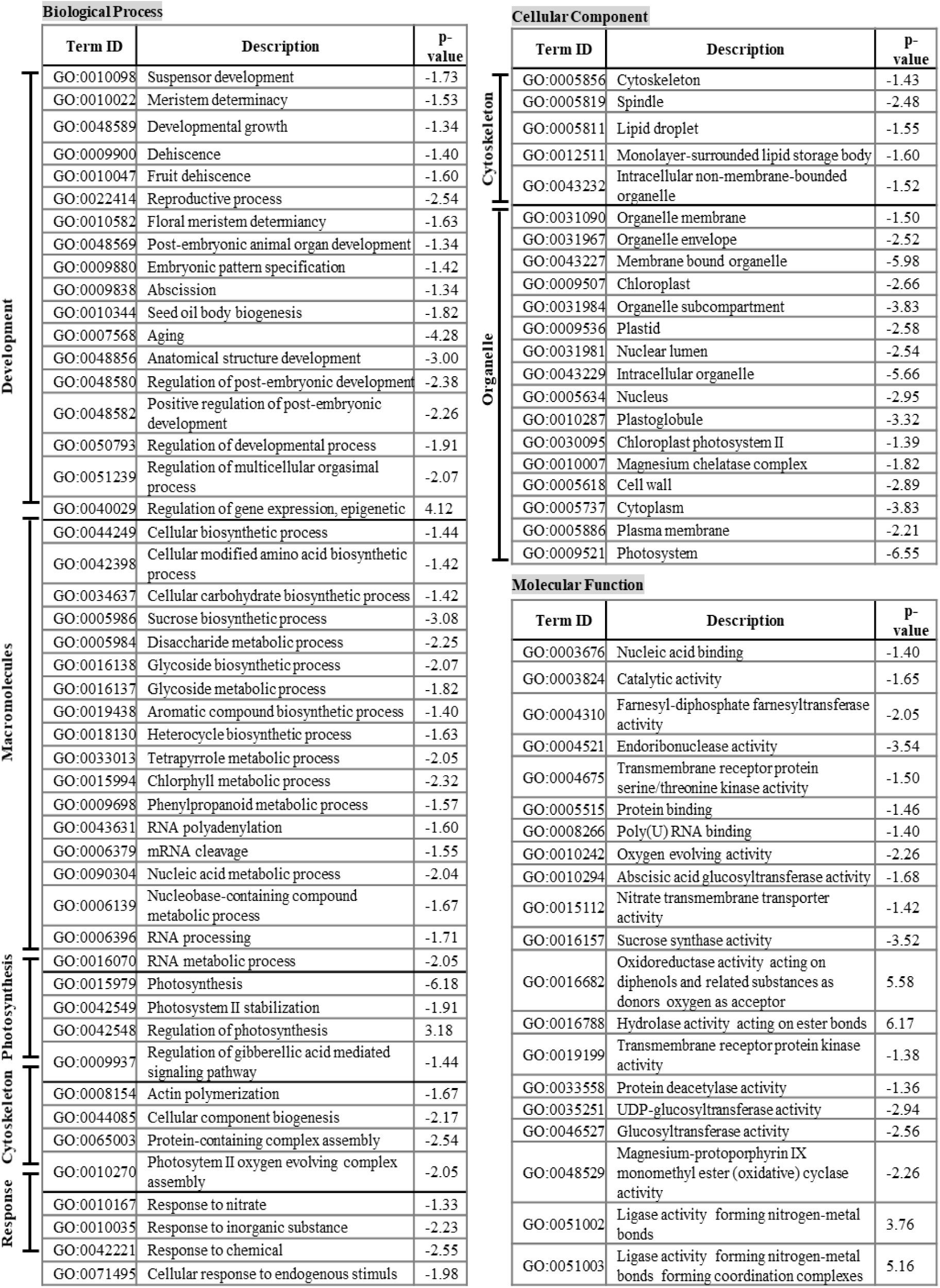
Enriched GO terms (biological processes, molecular function, cellular component) for upregulated DEGs in the calli of barley *cv*. GP

**Table 2B.**
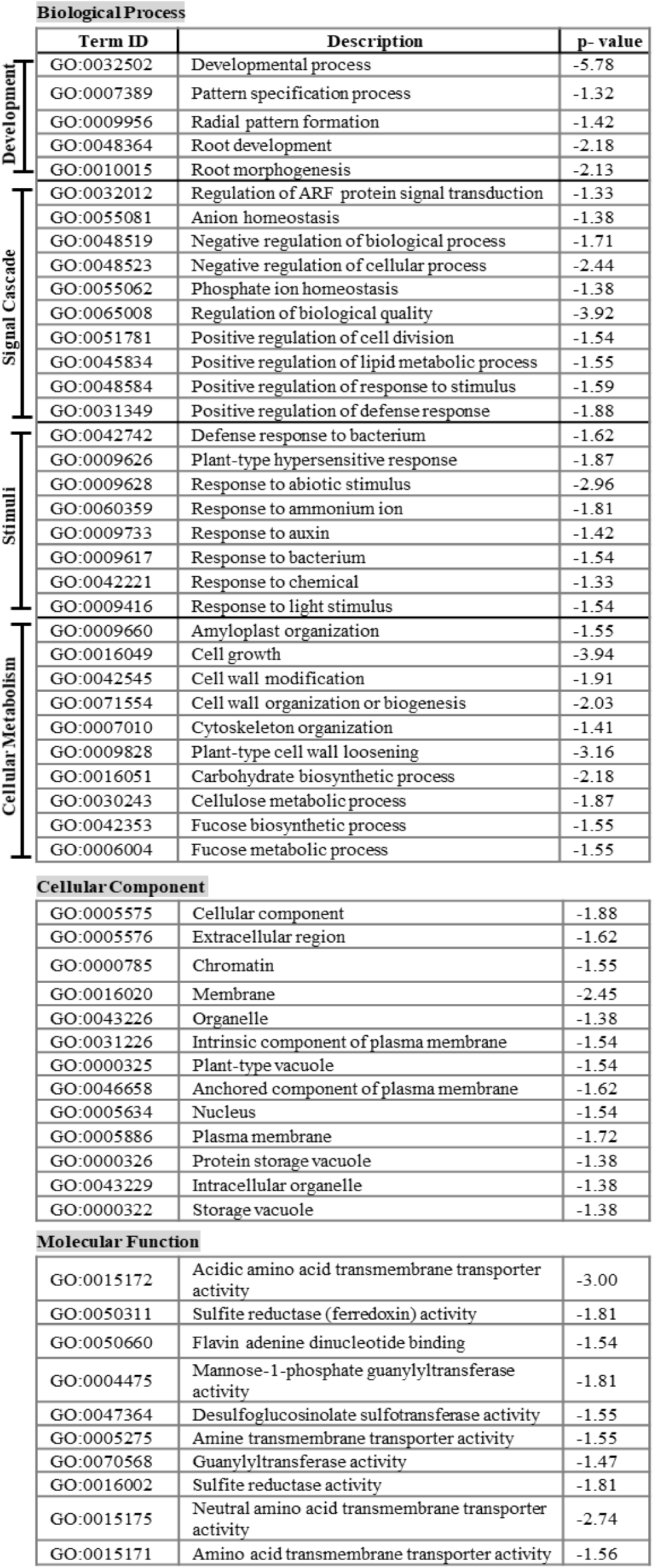
Enriched GO terms (biological processes, molecular function, cellular component) for downregulated DEGs in the calli of barley cv. GP

DEG analysis suggested that genes involved in the chlorophyll biosynthesis such as *Protoporphyrin IX Mg-chelatase subunit* (*HORVU2Hr1G023540*), *Magnesium-protoporphyrin IX monomethyl ester cyclase* (*HORVU3Hr1G039930*), and genes which regulates photosynthesis such as *chlorophyll a–b-binding protein* (*HORVU5Hr1G109250*), *photosystem I reaction center subunit X* (*HORVU3Hr1G009210*), *photosystem I reaction center subunit III* (*HORVU5Hr1G100140*), *photosystem I reaction center subunit VI* (*HORVU0Hr1G001490*), *photosystem I reaction center subunit N* (*HORVU2Hr1G019820*), *ribulose bisphosphate carboxylase small chain* (*HORVU5Hr1G051010*), *photosystem II protein D1* (*psbA*), *photosystem II reaction center Psb28 protein* (*psbD*) were significantly upregulated in the GP calli as compared to D91 (Fig. 5 and Supplementary Table S4). Transcripts related to carbohydrate metabolism were also found to be upregulated in GP, such as *fructose-bisphosphate aldolase* (*HORVU4Hr1G019570*), *sucrose synthase* (*HORVU7Hr1G007220*), *glyceraldehyde-3-phosphate dehydrogenase* (*HORVU4Hr1G082700*) (Fig. 5 and Supplementary Table S4). Besides, a large amount of the DEGs were involved in oxidoreductase activity, including several ROS (Reactive Oxygen Species)-related genes such as *iron/ascorbate-dependent oxidoreductase* (*HORVU5Hr1G065620*), *peroxidase* (*HORVU2Hr1G018480*), and *glutathione-S-transferase* (*HORVU6Hr1G090560*) were also found to be upregulated in *cv.* GP (Fig. 5 and Supplementary Table S4).

**Fig. 5.**
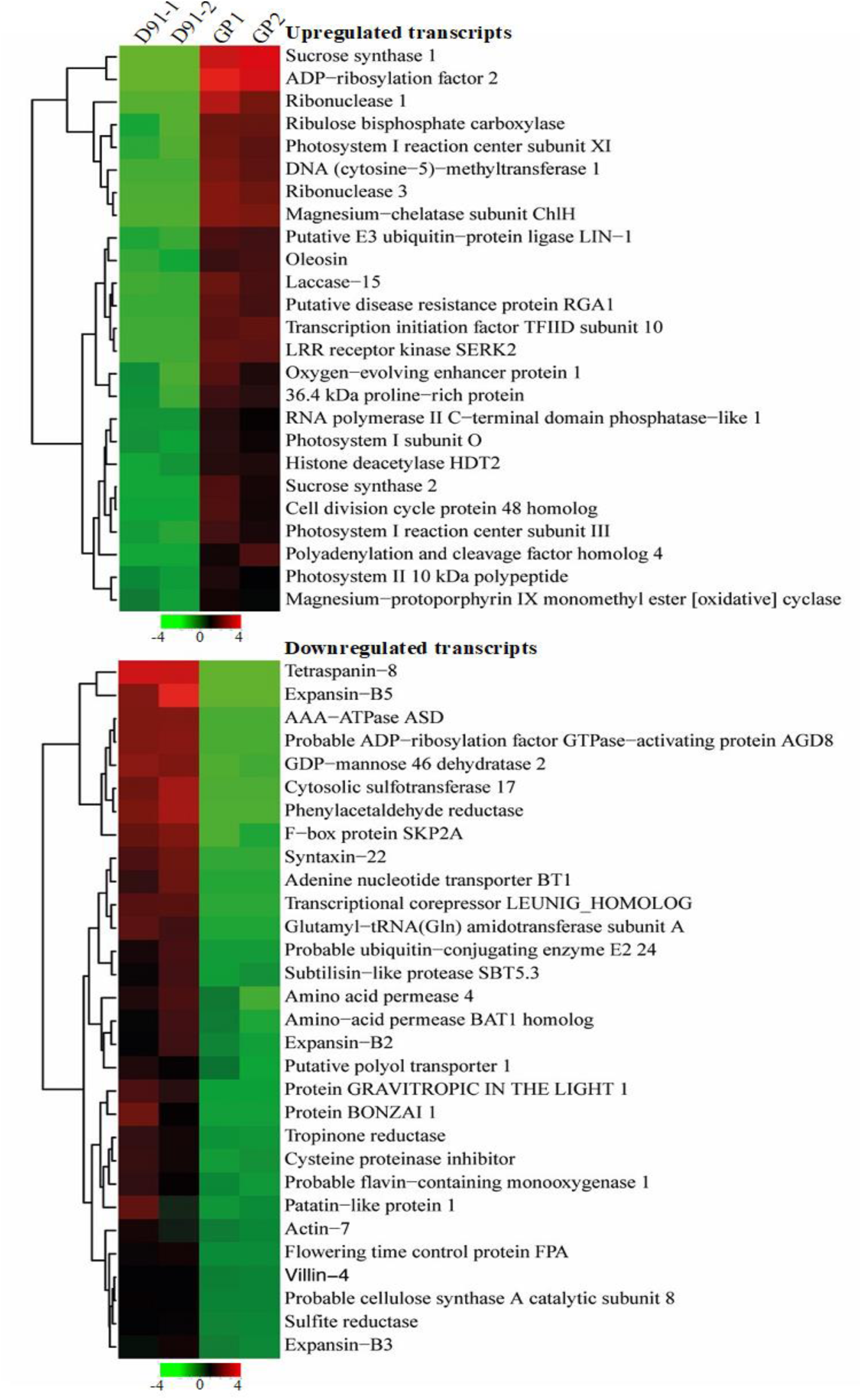
Heatmap showing the hierarchical clustering of the upregulated and downregulated DEGs in the calli of barley *cv.* GP. Green color represents downregulation and red color represent relatively high expression. TMM normalized FPKM values were used to construct the heatmaps. See color legend for expression levels.

### qRT-PCR validation of DEGs

qRT-PCR was performed on a sub-set of DEGs to validate the expression profile obtained using RNA-seq. Seven DEGs were selected for qRT-PCR, i.e., *HORVU6Hr1G058000* (*WAX2*), *HORVU1Hr1G007720 (Cytokinin*-O-glycosyltransferase2), *HORVU5Hr1G065620* (*Leucoanthocyanidin dioxygenase*), *HORVU2Hr1G015980* (*Nicotinamine synthase*), *HORVU7Hr1G113270* (*Endochitinase* A-*like*) and *HORVU2Hr1G107480* (non-*specific lipid*- transfer *protein*, *LTP*) (Fig. 6). qRT-PCR expression profile of the selected DEGs were in accordance with the transcriptome profile. Moreover, these results indicate that the genes are already differentially upregulated in *cv.* GP during the 3 days’ light period, as compared to D91 during early stages.

**Fig. 6.**
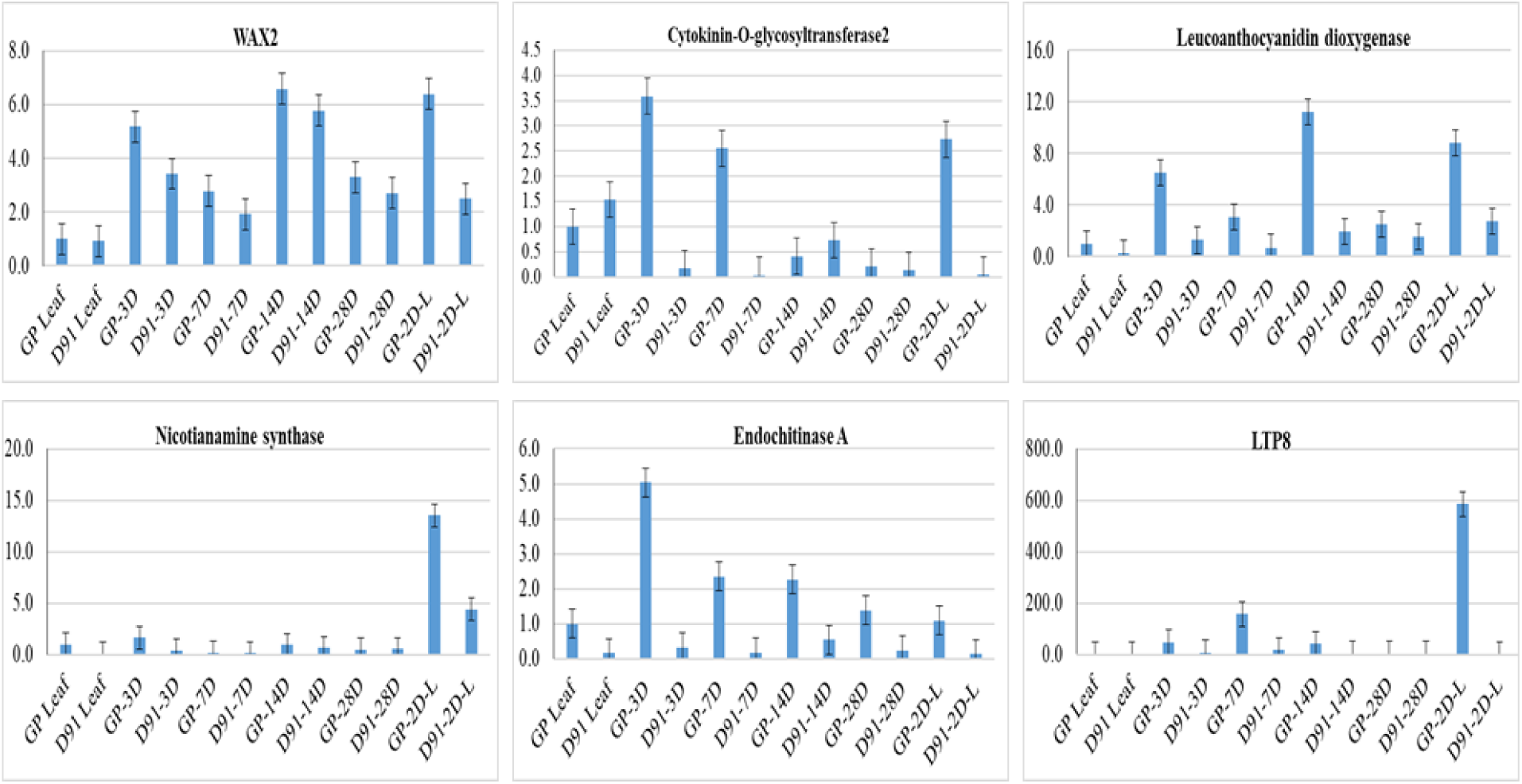
qRT-PCR analysis of 6 genes found to be upregulated in RNA-seq data. y-axis represent the change in expression level of these genes in terms of relative fold change, x-axis represents different time stages of GP and D91 calli. Relative fold change was calculated using 2^^-ddCt^ method.

### Subcellular localization of DEGs

Gene annotations were given to transcripts based on the similarity to the known genes. To further cross-check the analyzed RNA-seq data and the gene annotations, subcellular localization of selected transcripts was done. To confirm the subcellular localization of the selected DEGs, the corresponding full-length coding sequences were cloned into GFP fused destination vectors, *i.e., pB7FWG2*. As shown in Fig. 7, WAX2, Lipid transfer protein (LTP) were shown to target cytoplasm. Cytokinin-O-glycosyltransferase2 and Leucoanthocyanidin dioxygenase were localized in both cytoplasm and nucleus. Phosphogluconate dehydrogenase, Nicotinamide synthase, and chlorophyll a/b binding protein were localized in chloroplast. Endochitinase A was found to be localized in both cytoplasm and chloroplast. These results indicate an important role of chloroplasts in somatic embryogenesis.

**Fig. 7.**
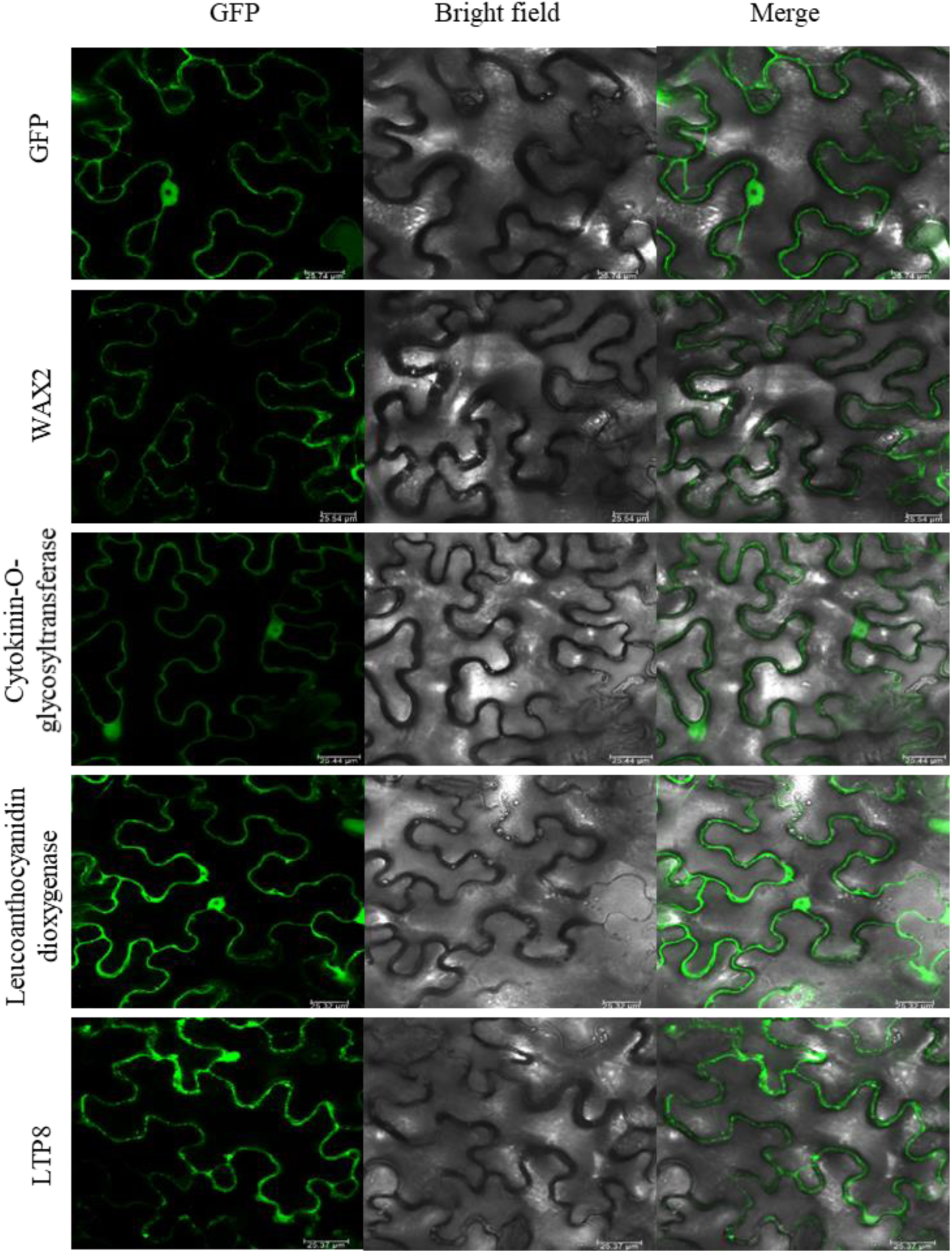
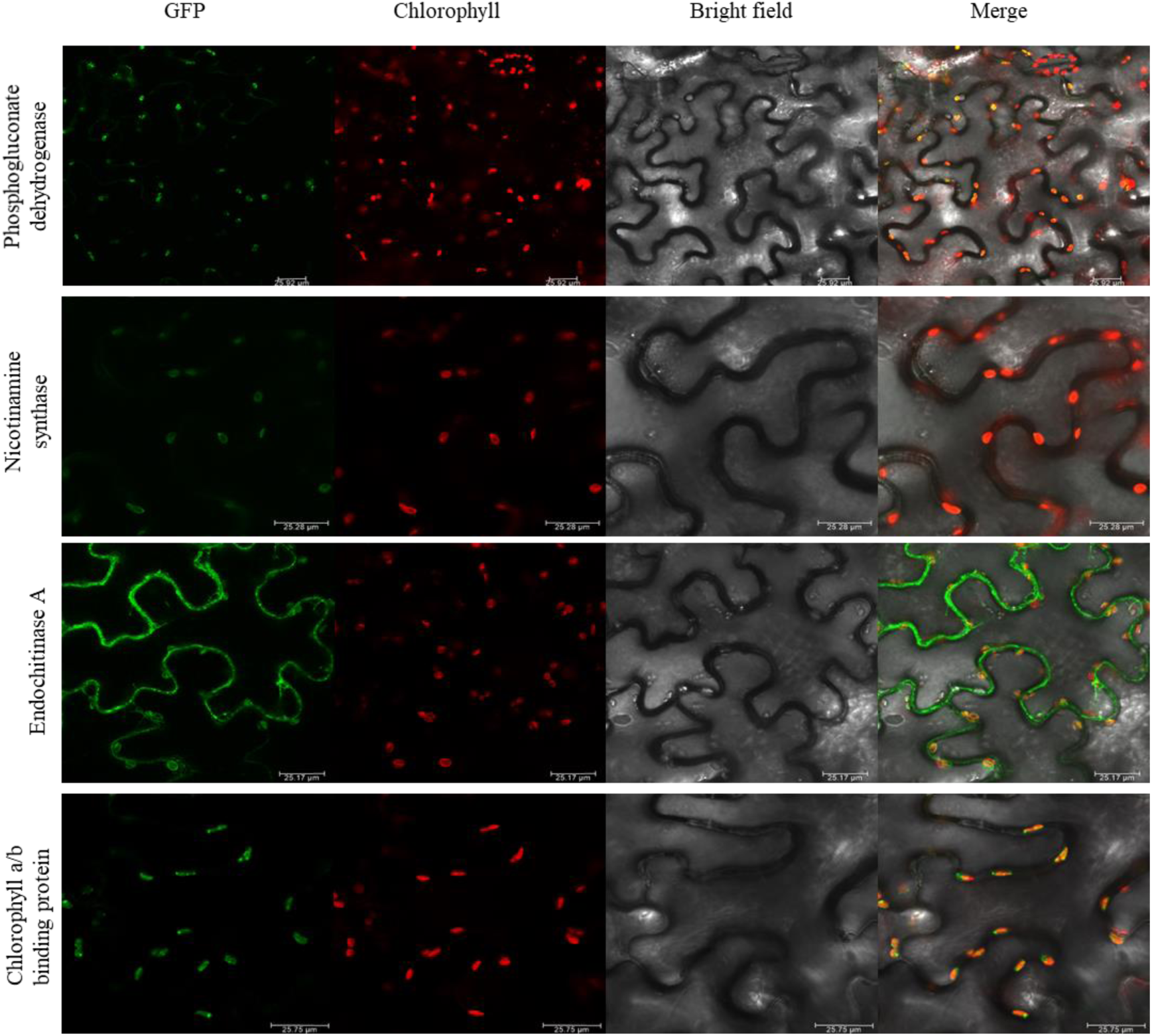
Subcellular localization of selected DEGs: C-terminal GFP fused construct. For GFP, WAX2, Cytokinin-O-glycosyltransferase2, Leucoanthocyanidin dioxygenase and LTP8, GFP emission is shown in the left panel, DIC in center panel and overlay in left panel. For phosphogluconate dehydrogenase, nicotinamine synthase, endochitinase A and chlorophyll a/b binding protein, GFP in shown in left panel, chlorophyll autofluorescence is shown in 2^nd^ panel, DIC is shown in 3^rd^ panel and overlay is shown in last panel.

### GP-D91 proteome profile

We performed proteomics analysis in two barley cultivars (GP and D91) using LC-MSMS. We identified 3062 protein groups and 16989 peptide groups (Supplementary Table S5), where 108 proteins groups were unique to GP and 82 protein groups were unique to D91 proteome (Fig. 8A). PCA and volcano plot analysis was performed to assess the variations and significance among the barley *cvs*. GP and D91 (Fig. 8B-C). The PC1 and PC2 represented 88.6% and 4.1%, total variance across the two barley cultivars, respectively (Fig. 8B-C). Moreover, 1586 DAPs were identified using t-test and Z-scaling method (Supplementary Table S6) and after removing the unannotated hits, 1005 protein groups were left which were further used for downstream analysis.

**Fig. 8.**
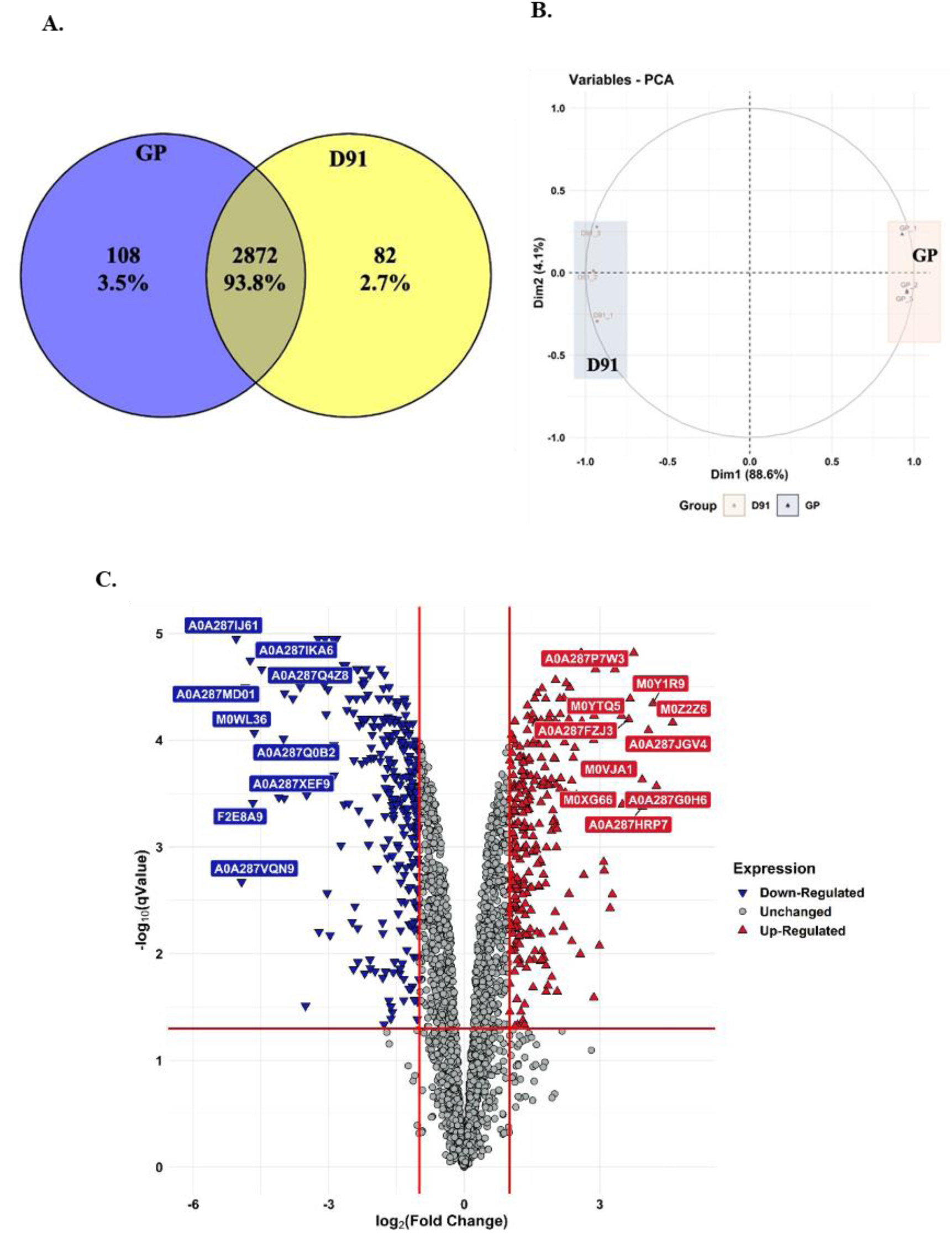
Comparison of proteome profile between GP and D91. (A) Venn diagram showing the differentially abundant proteins (DAPs). (B) principal component analysis (PCA) (C) volcano plot in barley *cv.* GP and *cv.* D91.

GO enrichment analysis revealed that categories like photosystem, photosynthesis, thylakoid, chloroplast-thylakoid membrane, photosynthetic membrane, plastid-thylakoid membrane, and cellular metabolic process were highly enriched in the upregulated DAPs of *cv.* GP (Fig. 9A) whereas categories such as carboxylic acid metabolic process, cellular amino acid biosynthetic process, and oxoacid metabolic process were enriched in the downregulated DEPs of barley cultivar GP (Fig. 9B). Additionally, pathway analysis exhibited different patterns for both the up- and down-regulated DEPs of GP-D91 proteome. Upregulated DEPs (Fig. 10A) in the *cv.* GP suggests the interplay of photosystem components (photosystem I and photosystem II), which regulates the plastid development and photosynthesis. On the contrary, downregulated DAPs in the *cv.* GP showed the network of catabolic processes (Fig. 10B). Proteins such as ribulose bisphosphate carboxylase (rubisco), phytoene dehydrogenase, cytochrome c domain-containing protein, cytochrome-f, ATP synthase, photosystem I iron-sulfur center, photosystem I P700 chlorophyll a, chlorophyll a-b binding protein, photosystem II reaction center, psbp domain- containing protein, and DNA replication licensing factor MCM7, MCM2, MCM6, and Malate dehydrogenase were found to be differentially abundant in the *cv.* GP (Supplementary Table S6) whereas proteins like acetyl-coenzyme A synthetase, acyl-coenzyme A oxidase, fatty acyl-CoA reductase, ferredoxin-NADP reductase, and ferritin were abundant in the *cv.* D91 (Supplementary Table S6). MS analysis suggests a higher abundance of protein related to chloroplast biosynthesis and photosystems in the GP whereas D91 lacks the proteins related to photosynthesis and chloroplast biosynthesis.

**Fig. 9.**
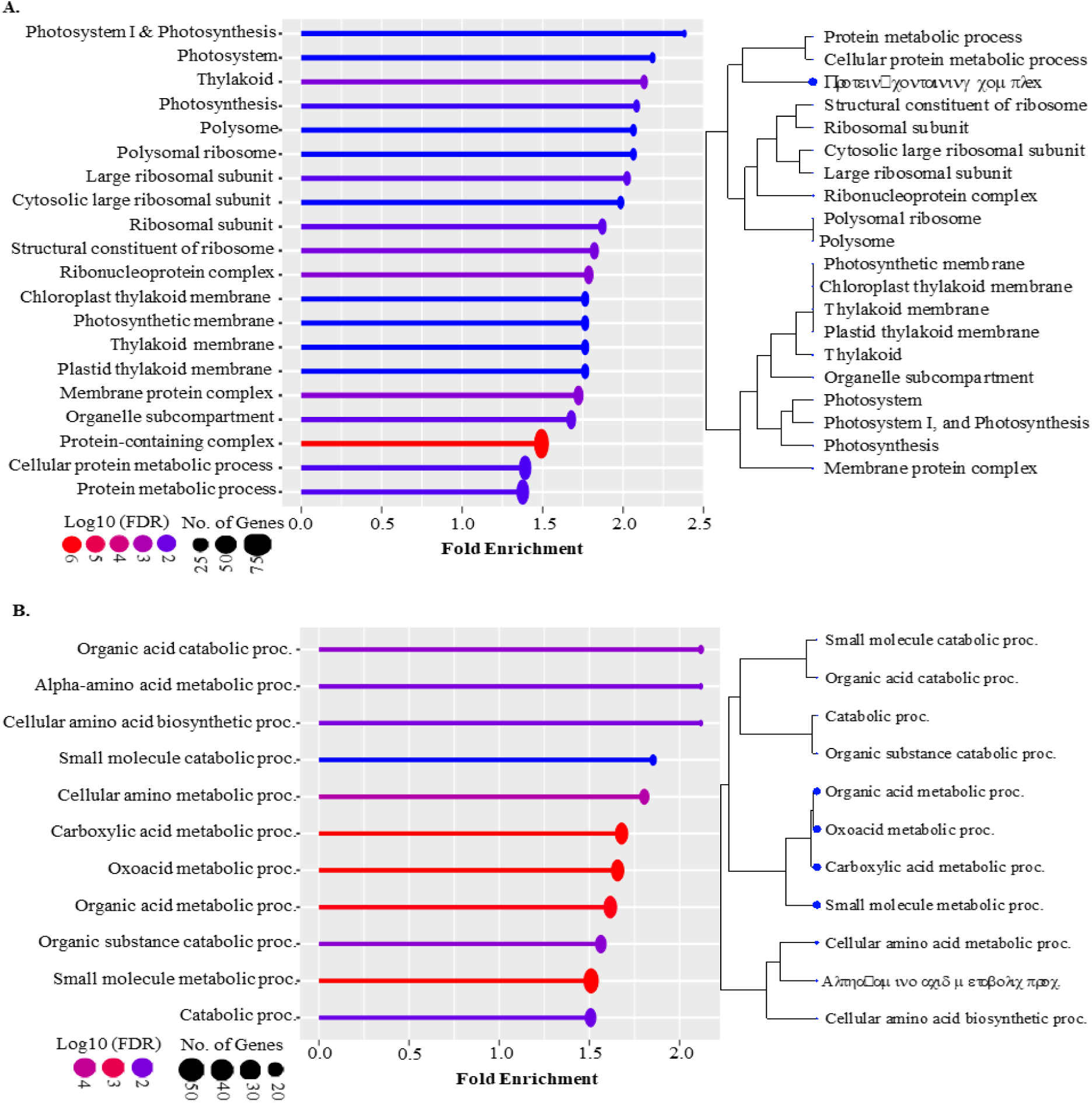
Bar charts and cladograms showing the enriched GO terms in barley *cvs.* A.) GP-up B.) GP-down.

**Fig. 10.**
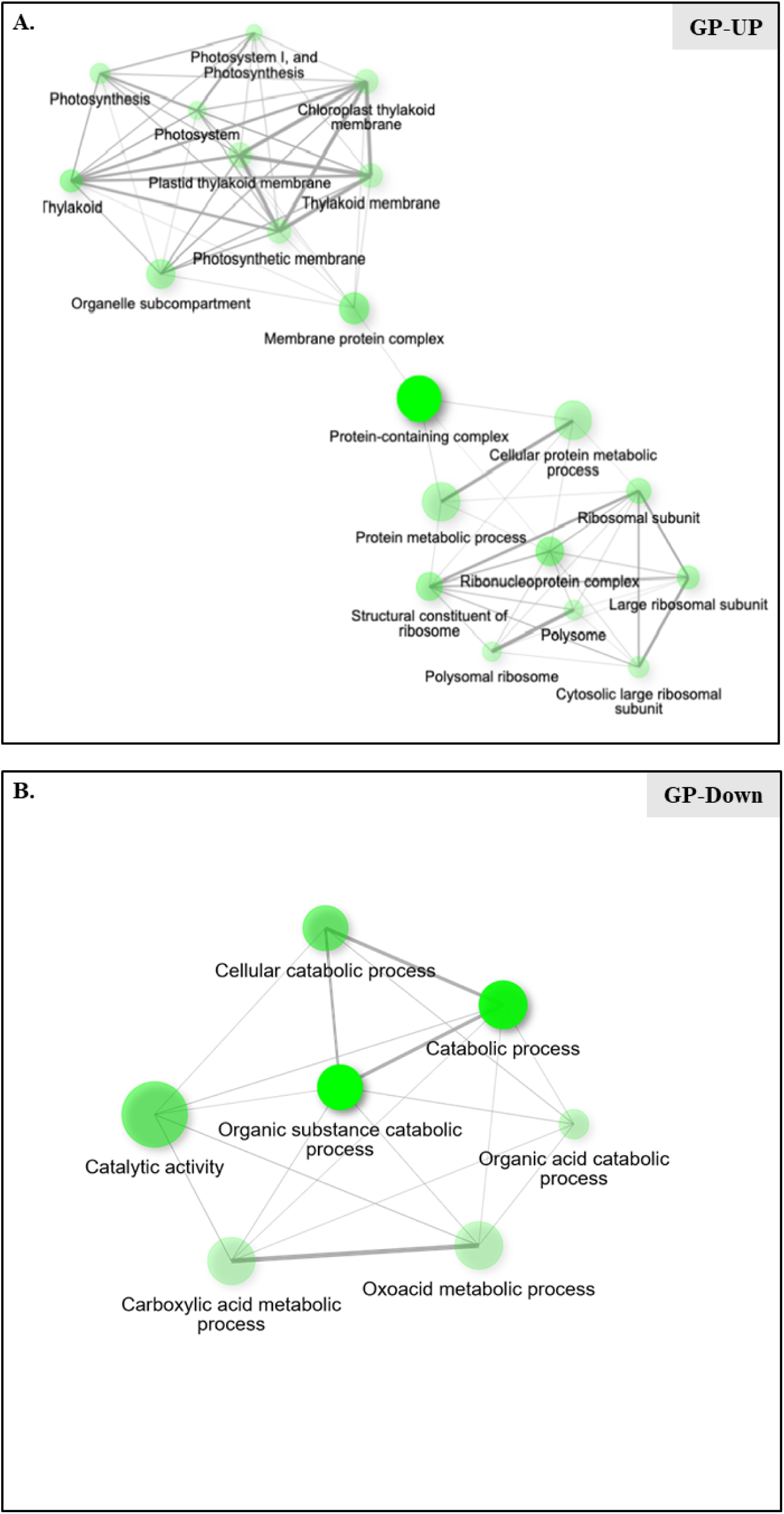
Network analysis showing the major pathways enriched in barley *cvs.* A.) GP-UP B.) GP-DOWN using web server shinyGO.

### Reactive Oxygen Species is important for somatic embryogenesis

ROS is produced as a normal product of plant cellular metabolism (Sharma et al., 2012) which also known to facilitates somatic embryogenesis (Zhou et al., 2016). Our RNA-Seq data also showed presence of pathways such as oxidation-reduction, and photosynthesis related to ROS in GP as compared to D91. Thus, we next checked the presence of free oxygen radicals in calli of both GP and D91 by NBT staining method (Wohlgemuth et al., 2002) and results showed that the characteristic blue colour of the GP calli was more intense than that of the D91 calli (Fig. 11A), indicating that more ROS was accumulated in GP calli compared with the D91 calli. Additionally, quantitative estimation of formazan precipitate in the calli was done using NBT staining. GP calli showed higher concentration of formazan at both stages *i.e.,* 30-days calli on BCIM and 30 days + 3 days calli on BRM (Fig. 11B).

**Fig. 11.**
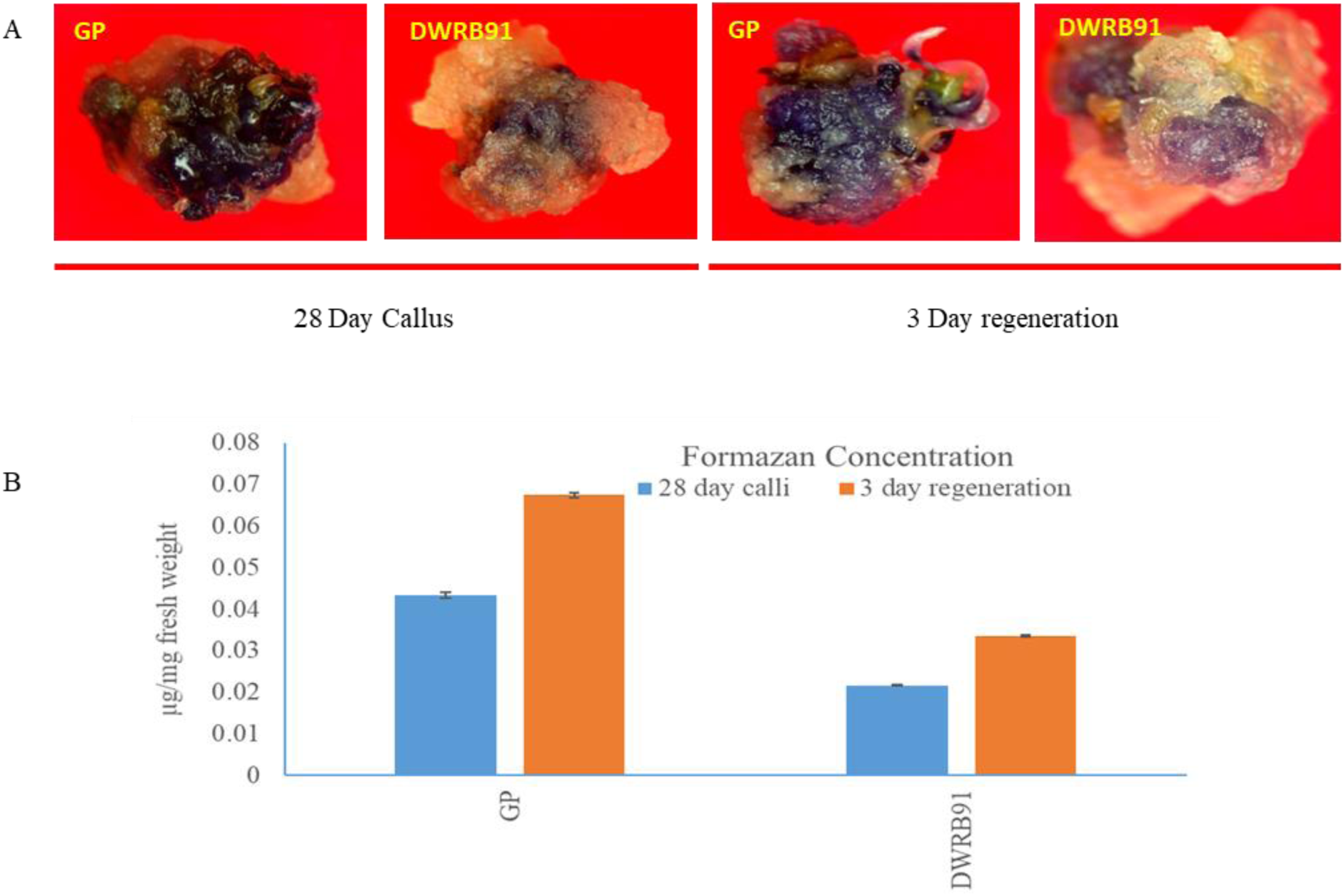
NBT staining and formazan quantification. (A) Detection of reactive oxygen species by NBT staining in the callus of Golden Promise and DWRB91 at 28-day callus stage and 28 days+ 3days regeneration stage. (B) Quantification of formazan in Golden Promise and DWRB91 callus.

### Metabolic profiling reveals alterations in primary and secondary metabolites of Golden Promise and DWRB91 callus

Callus of GP and D91 (28 days + 3 days) were used for comparative metabolomics. A total of 36 differentially accumulating metabolites were identified by GC-MS metabolite profiling of GP and D91 callus metabolites, which includes amino acids, organic acids, secondary metabolites, and sugars. Total ion chromatograms of GC-MS run are shown in Figure S2. Metabolite details, their retention time and quantification ions (Q-ions) are presented in Supplementary Table S7. The one-way ANOVA p-value < 0.05 was used to compare the metabolites. Principal component analysis (PCA) was used to visualize the discriminating differences between both callus types (Fig. 12A). The first and second PCs of the PCA score plot represented 79% (PC1) and 21% (PC2) of the total variance of the samples, which revealed the important cleavage between embryogenic calli of GP and non-embryogenic calli of D91. To check the contribution of differential metabolites in GP and D91 callus, the PLS-DA was conducted. The 2D score plots obtained from the PLS-DA test showed that the metabolites of GP calli were distinctively different from D91, indicating an altered state of metabolite levels in the callus of these cultivars (Fig. 12B). The component 1 and component 2 of PLS-DA accounted for 78.8 % and 21.2 % variance among samples, respectively. A VIP score plot that displays metabolites with VIP scores greater than the cut-off value of 1 was used to further assess the relevance of the metabolites (Fig. 12C). Twenty-two significantly differential marker metabolites were detected based on VIP scores (Fig. 12C). The heatmap for all the differentially accumulating metabolites used for the PCA analyses is shown in Fig. 12F. The heatmap offered excellent separation of the metabolite trend between the GP and D91 calli samples. Metabolites related to carbohydrate metabolism, such as sucrose, fructose, glucose were found to be highly accumulated in GP. Subsequently, we observed that proline and lysine were also significantly higher in GP callus. Also, metabolites such as p-coumaric acid, cinnamic acid, protocatechuic acid, caffeic acid and shikimic acid which act as the precursors of secondary metabolites and also involved in lignification of cell walls, showed enhanced accumulation in GP calli as compared to D91 calli.

**Fig. 12.**
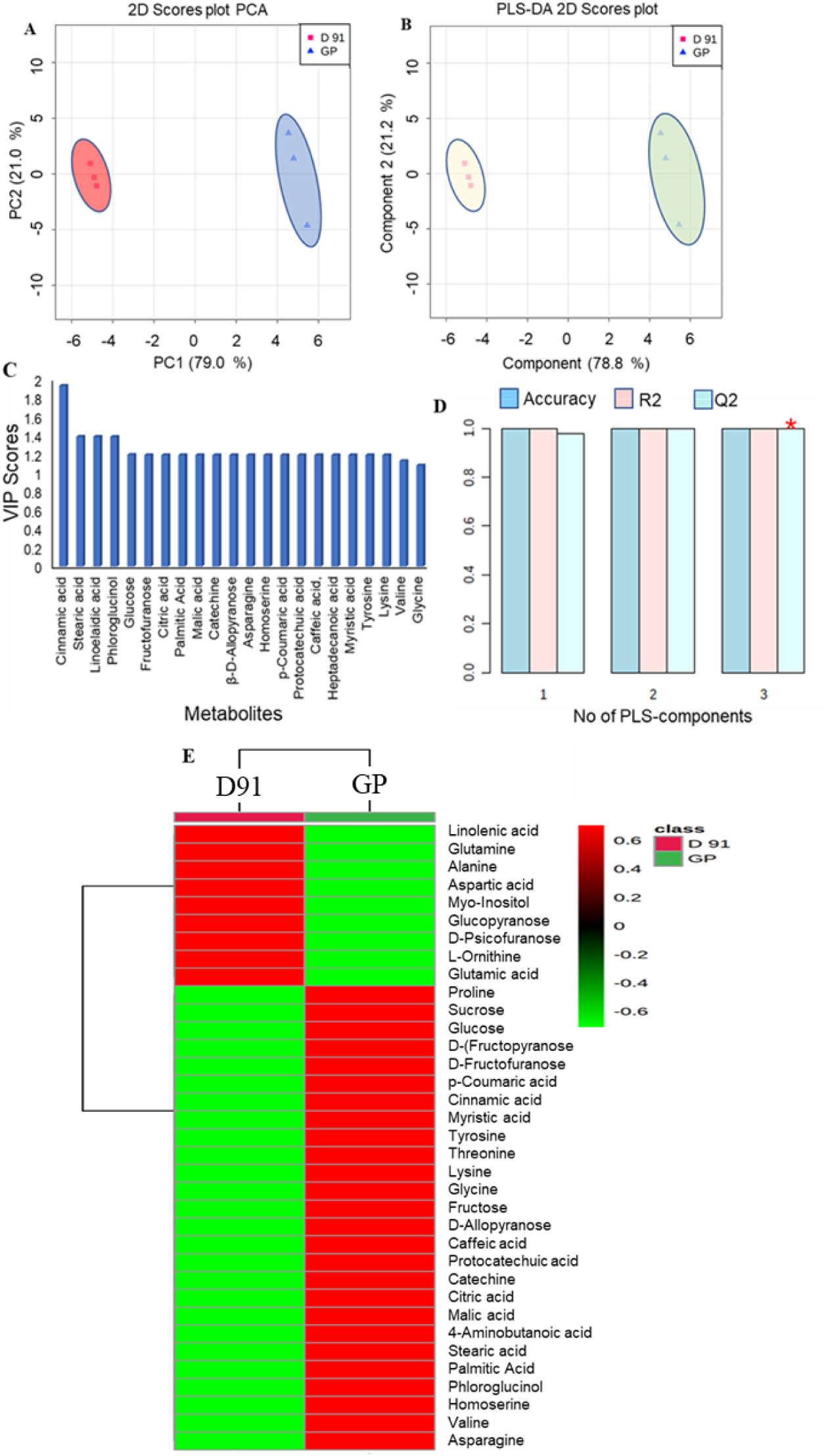
Variation in the metabolite profile between calli of GP and D91. (A) PCA score plot showing distinct clustering of two sample types; (B) PLS-DA scores plot showing separation of samples; (C) variable importance in project (VIP) scores of top fifteen volatiles which substantially contributed in discriminating different postharvest storage time points (D) Performance of PLS model with respect to different principal component numbers; (E) Heatmap of all metabolites between two sample types. R2: cumulative explained variation and Q2 predicted variation.

The identified differential metabolite markers were used for pathway analyses by Metaboanalyst 3.0, which showed alteration in sugar metabolism, tricarboxylic acid cycle (TCA) metabolism, fatty acid biosynthesis, secondary metabolite biosynthesis, glycine/serine metabolism and phenylpropanoid metabolism (Fig. 13, 14). It is noteworthy here that these metabolites are synthesized in plastids/chloroplasts, and it is nicely synced with the differential gene expression of respective genes as revealed by transcriptomics and proteomics (Fig. 6).

**Fig. 13.**
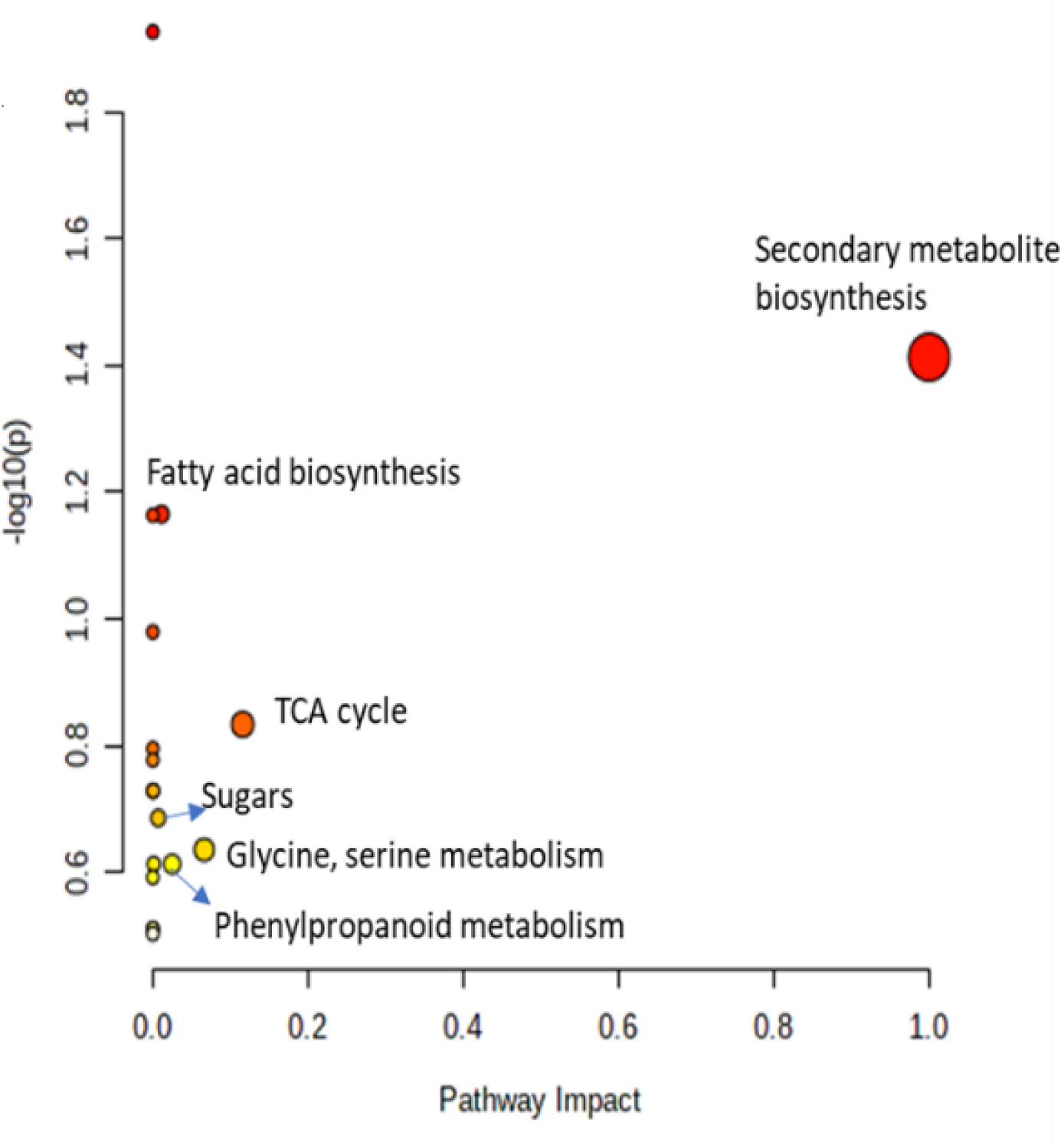
Metabolomic Pathway Analysis (MetPA) plot created by MetaboAnalyst (5.0) using Rice library. In the panel, the matched pathways are presented as circles where the color intensity of the circle is based on p-value significance (darker circle colors denote higher significant changes of metabolites in the respective pathway). The size of each circle corresponds to the pathway impact score. The pathways having high impact scores and significance levels are annotated.

**Fig. 14.**
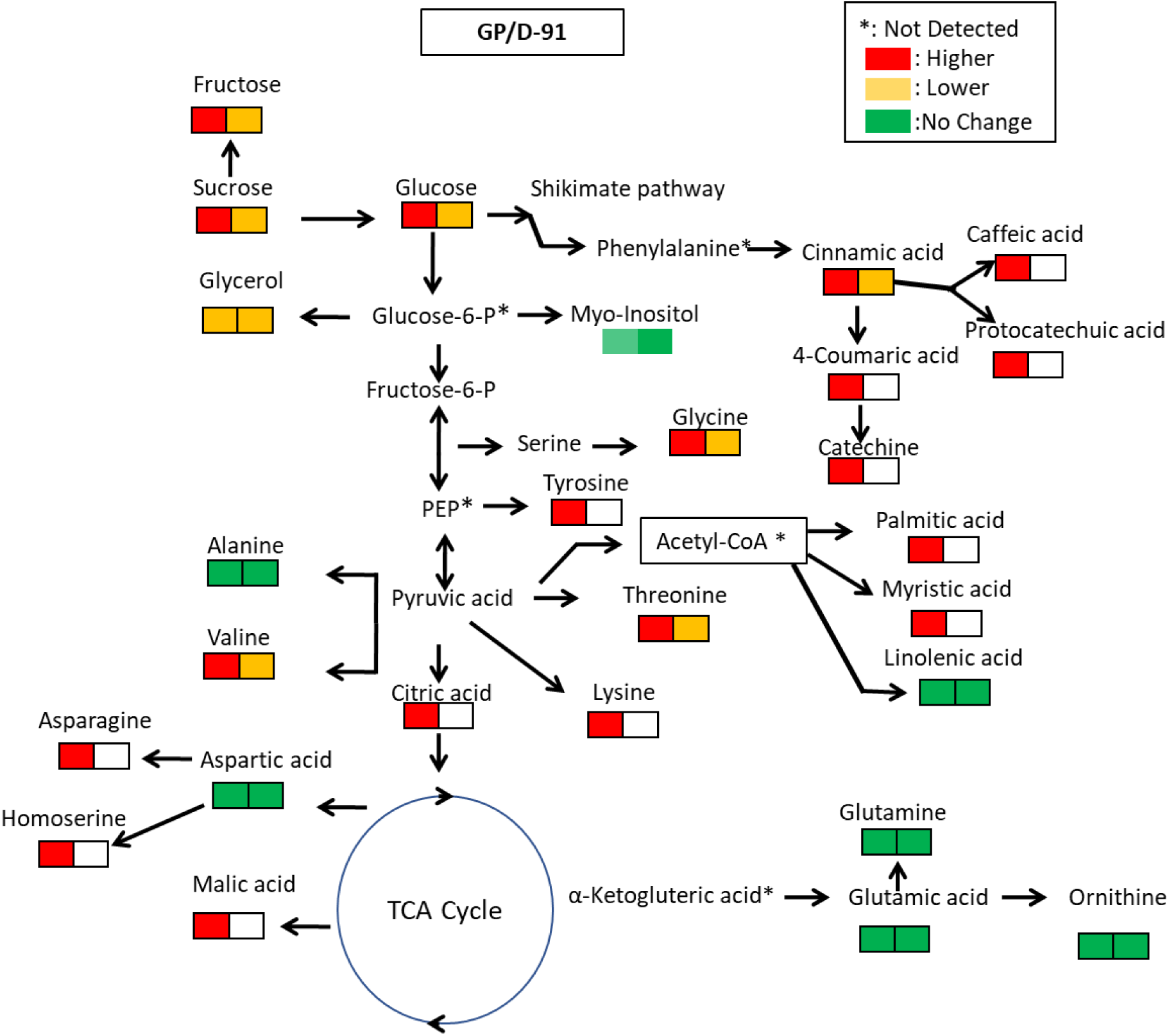
Pathway network showing the major metabolic changes detected in calli of GP and D91. * = Metabolites incorporated in the pathway based on prior pathway information (not detected in the present analyses)

### Lignin biosynthesis is important during somatic embryogenesis

Lignification provides strength to cell wall which is also related to differentiation processes. Comparative metabolomics revealed a major class of metabolites involved in lignification process. To investigate a potential relationship between *in vitro* regeneration and lignin biosynthesis, we examined lignin accumulation in the 4-weeks-old calli of both GP and D91. There is significantly more lignin content in the calli of GP than D91 (Fig. 15A). This is further evident from morphological characterization through phloroglucinol–HCl staining of the calli revealed significant lignification in GP callus (pinkish red stain), especially in the areas of somatic embryos as compared to D91 callus (Fig. 15B).

**Fig. 15.**
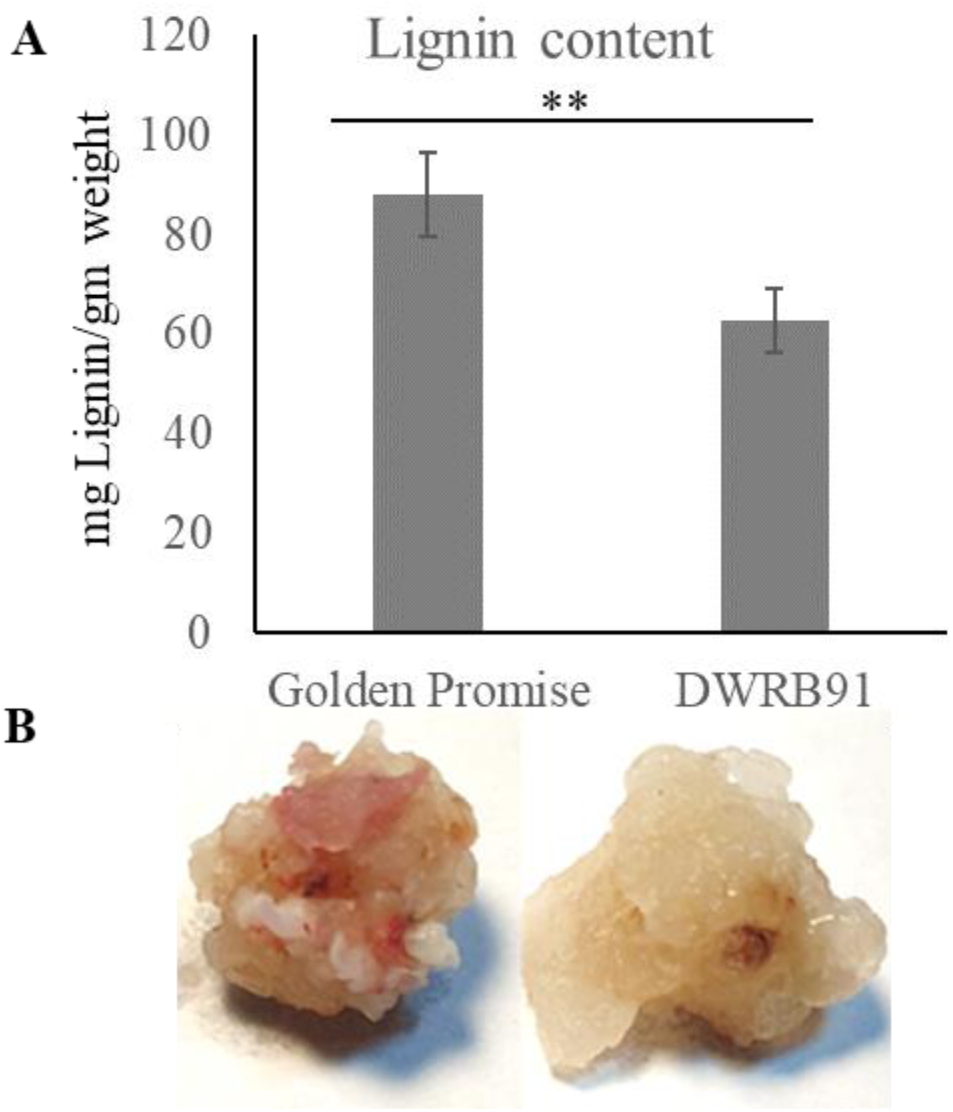
Detection of lignin by phloroglucinol staining in the callus. **A).** Quantification of lignin in Golden Promise and DWRB91 callus at 28 days + 3days regeneration stage. B). Detection of lignin by phloroglucinol staining in the callus of Golden Promise and DWRB91 at 28 days+ 3days regeneration stage.

## Discussion

Most of crop plant species have low regeneration efficiency under *in vitro* conditions, as the *in vitro* regeneration efficiency of plants greatly depends on the formation of embryogenic callus which can differentiate into new plants. The existing regeneration methods mainly use mature and immature embryos as starting material and are highly genotype-dependent for most of the crop plants. Barley is the fourth most important cereal crops worldwide after rice, wheat, and maize. However, barley genetic improvement is highly limited mainly due to genotype dependency for *in vitro* regeneration. GP is the model cultivar used for barley transformation which is subject to seasonal constraints. In contrast, Indian cultivars are highly recalcitrant to *in vitro* regeneration via tissue culture and low potency of embryogenesis. Considering the above factors, we have taken barley as the model crop for understanding the mechanism of *in vitro* regeneration in temperate cereals. In this study, we explored morphological, global transcriptional, proteome, and metabolomics changes during regeneration of mature embryo-derived callus and identified some potential factors that might contribute to the differential responses of the two types of cultivars.

Callus induction plays a key role in the formation of somatic embryos. This study showed morphological analysis through SEM to analyze callus in the barley *cvs.* GP and D91. This study showed morphological variations between embryogenic calli of GP and non-embryogenic calli of D91. Similar results have been reported in different crops including *Panicum virgatum*, *Oryza sativa*, *Stylosanthes spp*. and *Cenchrus ciliaris* (Burris et al., 2009; Binte Mostafiz and Wagiran, 2018; Narciso and Hattori, 2010; Bevitori et al., 2014; Abe et al., 1986; Andi et al., 1992; Kumar and Chandra, 2010; Kumar et al., 2015). Also, the relative water content of GP calli was found to be significantly less than D91, which shows that low relative water content is an important factor of somatic embryogenesis and regeneration. Previous reports demonstrated that out of 3 callus types of ravenna grass, calli with lowest water content showed highest callus proliferation, whereas the calli with highest water content showed lowest proliferation and regeneration (Shimomae et al.; 2013).

Transcriptomic studies of rice and maize led to the identification of important transcription factors such as *WOX5*, *BBM*, *WUSCHEL*, *LEC1*, *LEC2*, and *WUS2* which regulates somatic embryogenesis in recalcitrant genotypes when expressed ectopically, but still the frequency of somatic embryo formation is low (Richards et al., 2015; Boutilier et al., 2002; Sinohara et al., 2015; Florez et al., 2015; Horstman et al., 2017; Lowe et al., 2016). The process of callus maturation is still poorly explored and there is not much known about the molecular changes which regulates it. We compared the global gene expression of calli of two different cultivars of barley during critical developmental time point to identify the genes which are essential for cellular reprogramming and redifferentiation. GO analysis suggested the processes related to photosynthesis, carbohydrate metabolism, chlorophyll biosynthesis and ROS homeostasis were significantly enriched in the *cv.* GP. Chlorophyll biosynthetic genes like *Protoporphyrin IX Mg-chelatase subunit*, *Magnesium-protoporphyrin IX monomethyl ester cyclase* along with Chlorophyll a/b binding protein were upregulated in *cv.* GP and have been found to regulate chlorophyll accumulation in mature embryos and photosystems PSI and PSII in chloroplast. Similar results were reported by Gulzar et al. (2021) in *Catharanthus roseus* (L.) by analyzing protein profiles of cotyledonary embryos while attaining maturity. Genes belonging to carbohydrate metabolism were also found to be upregulated in GP which suggests the need of energy for maturation of embryos and importance of plastids/chloroplasts in the source-sink relationship. Moreover, proteome profiling corroborates the fact that genotype GP maintains the source-sink relationship in an energy efficient manner as comparison to genotype D91. Comparative metabolomics of GP and D91 calli further confirmed the higher accumulation of sugars such as sucrose, fructose and glucose. The accumulation of chlorophyll along with increased carbohydrate metabolism indicated better photosynthetic ability of GP as compared to D91, further confirming the crucial role of plastids/chloroplasts in *in vitro* regeneration.

Furthermore, we showed that the oxidation-reduction process and ROS generation processes were significantly enriched in GP. The results suggested that ROS accumulation promotes somatic cell embryogenesis, leading to increase of regenerating plantlets. The result is similar to ROS accumulation due to reduced SOD1 activity in *OsLOL1* overexpressed embryogenic cotton, thereby leading to increase in somatic embryogenic capacity, which suggests that high ROS level facilitates somatic embryogenesis in cotton calli (Zhian et al., 2020). In another study, importance of ROS signalling in somatic embryogenesis was investigated in cotton by culturing the explants on medium supplemented with different concentrations of DPI (Diphenyleneiodonium; NAPDH oxidase inhibitor) (Ellis et al, 1988). The DPI treatment led to the retardation of the dedifferentiation process indicating that ROS was necessary for dedifferentiation during cotton somatic embryogenesis (Zhou et al, 2016). Environmental stresses also lead to ROS accumulation in order to promote somatic embryogenesis and plant regeneration (Rai and Shekhawat 2011; Rodríguez-Serrano et al. 2012). The above evidence suggests that high ROS level in callus promotes transition of somatic-to-embryogenic callus cells and plant regeneration.

Further, metabolite profiling shows clear distinction between embryogenic calli of GP and non-embryogenic calli of D91. In particular, GP showed higher accumulation of amino acids which suggests the need of amino acids for cell division and differentiation, ultimately leading to plant regeneration. In our study, proline, lysine, asparagine was found in high concentration in GP. Liang et al. (2013) discussed the role of proline metabolism in promoting embryo formation by stimulating cell-signaling pathways through increased formation of reactive oxygen species (ROS). Gerdakaneh et al. (2011) reported that proline induced maturation of SE in strawberry. Proline was also found to be accumulated in embryogenic calli of sugarcane (Neves et al., 2003).

Proline is also associated with various signal transduction pathways which results in improved cell signaling and reported to improve somatic embryogenesis by providing nutrition (Mujib and Samaj, 2005). Similarly, lysine was reported to be accumulated in higher concentration in EC of *Boesenbergia rotunda* (fingerroor ginger) (Ng et al., 2016). Pongotongkam et al. (2004) also reported increase in rice plantlet regeneration by exogenous application of lysine. We have also found that calli of GP accumulated significantly more amount of these amino acids in comparison to calli of D91.

Carbohydrates play an important role during plant growth, developmental and floral transition by acting as signal for gene expression (Eveland and Jackson 2012). As indicated by heat map correlation, GP showed higher accumulation of important sugars such as fructose, glucose and sucrose. This increased accumulation of sugars suggests the importance of sugars in cell proliferation and differentiation processes (Abdelsalam et al., 2021). Hudec et al. (2016) also studied the changes in carbohydrate profiles of Norway spruce embryogenesis and found increase in sucrose: hexose ratio during maturation.

Lignin is an essential component of plant secondary cell walls, which influence plant growth and differentiation (Dauwe et al., 2007). In our study, phenolic compounds mainly the key intermediaries for lignin biosynthesis such as p-coumaric acid, cinnamic acid and caffeic acid were also found to be accumulated in high concentration in calli of GP as compared to D91. Similar results were reported by Awanda et al. (2019) for accumulation of chlorogenic acids during embryo development in *C. arabica*. Our results suggest that these differentially accumulating metabolites might be contributing to the somatic embryogenesis of GP calli.

In conclusion our data reveals the important part plastids play in the regeneration capabilities in contrasting genotypes. The regeneration competent calli showed higher plastid/chloroplast activity in terms of transcriptomics, proteomics, and metabolomics. Although from appearance only, it seems that all types of genotypes make callus under auxin supplemented nutrient media, however, it is the metabolic activity at both primary and secondary level and mainly involving plastids which make the difference between genotypes showing regeneration capability, thereby making them target for genetic engineering. Further studies can be undertaken in the direction of studying de novo plastid origin between contrasting genotypes and studying some suitable plastid mutant for their efficiency in *in vitro* regeneration.

## Supplementary data

Figure S1. In vitro regeneration response of different barley genotypes

Figure S2. A representative total ion chromatogram (TIC) of GC-MS analyses of D-91 and GP variety. Peak details and their identification are given in Table S5.

Table S1. List of primers used in qRT-PCR expression analysis.

Table S2. List of primers used for cloning for sub-cellular localization.

Table. S3. Details of RNA Sequencing.

Table S4. List of differentially expressed genes.

Table S5. List of peptides identified using LC-MS/MS (LFQ) for barley cvs. GP1 and D91.

Table S6. List of differentially abundant proteins (DAPs) identified using Z-scaling method for barley *cvs*. GP and D91.

Table S7. Peak number details and compound names identified for significant metabolites between GP and D91.

## Abbreviations

GP: Golden Promise
D91: DWRB91
DEG: differentially expressed gene
ROS: Reactive oxygen species
DEP: differentially expressed proteins
SE: somatic embryogenesis
EC: embryogenic callus
NEC: non-embryogenic callus

## Acknowledgements

PS thanks MHRD for Junior and Senior research fellowships. We thank Genetic Resource Center, National Agriculture and Food Research Organization Japan for FL-cDNA used in subcellular localization.

## Author contributions

HC conceptualize the study, HC, PS, CC, SB, RV, SKM, RC, BW and AP performed experiments, HC, PS, CC, HG, DS analyzed the data, HC, PS and CC wrote the manuscript. All authors read and approved the submitted version.

## Data availability

Data generated/analyzed and supplementary information in this study is available with the published article. Raw data and files are available from the corresponding author on request.

